# Mechanism-guided Discovery of Fluoroacetate Dehalogenase (FAcD) Variants for Enzymatic PFAS Defluorination

**DOI:** 10.64898/2026.10.11.758284

**Authors:** Shaban Ahmad, Asal Forouzandeh, Benedetta Togni, Jonas Borup Randløv, Michael Ogden, Tue Kjærgaard Nielsen

## Abstract

Per- and polyfluoroalkyl substances (PFAS) are persistent anthropogenic “forever chemicals” whose strong C-F bonds (∼460–533 kJ mol⁻¹) resist defluorination, and the recast EU Drinking Water Directive sets parametric values of 0.10 µg L⁻¹ for the sum of 20 specified PFAS and 0.50 µg L⁻¹ for total PFAS. Here, we present **DeFluorX**, a mechanism-first pipeline that uses Boltz-2 to co-fold 2,150 bacterial fluoroacetate dehalogenase (FAcD) variants with 26 PFAS and fluoroacetate, generating uniformly high model confidence that tracked catalytic competence poorly (ρ = +0.08). Thus, the complexes were ranked and tiered by S_N_2-defluorination geometry based on the known mechanism of control of FAcDs RPA1163 with fluoroacetate: Oδ···Cα distance and Oδ-Cα-F angle; relay; carboxylate clamp; F-stabiliser; F-cradle; and penalties for scissile bond strength, backside occlusion, β-fluorination and bond multiplicity. The tiering identified eight complexes (med ∠ 173.2°, distance 2.91 Å) with a high probability of degradation, including Alphaproteobacterial enzymes, with mono-, di- and trifluoroacetate attacking at 172–178° versus 163° for the control. We performed WaterMap, restrained 1 µs Desmond MD, MM-GBSA and scans of the three WaterMap-aware best MD frames per complex using B3LYP/6-31G** QM/MM. All 12 frames exhibited product-end C-F cleavage with barriers ranging from 1.1 to 11.8 kcal mol⁻¹, and the departing fluoride carried −0.43 to −0.63 e. The Laboratory validation is ongoing.

## 1. Introduction

Per- and polyfluoroalkyl substances (PFAS), widely referred to as "forever chemicals", are a structurally diverse class of anthropogenic compounds that persist in water and biota, and their association with adverse health outcomes has prompted regulatory action ^1,2^. The EU Drinking Water Directive sets regulatory limits of 0.10 and 0.50 µg L⁻¹ for the sum of 20 listed substances and for PFAS total, respectively ^3^. Chemical and photocatalytic routes to complete mineralisation exist and have advanced quickly ^4,5^, but they operate far from the dilute, ambient conditions of contaminated groundwater. Biological defluorination is attractive because it does not require them, but the lack of known enzymes that effectively degrade PFAS is a major obstacle to microbial biotransformation ^6,7^.

Bacterial fluoroacetate dehalogenase (FAcD, EC 3.8.1.3) is the best-characterised enzyme class that hydrolytically cleaves a C-F bond; it converts fluoroacetate (FA) to glycolate and releases fluoride under ambient conditions ^8^. The C-F bond it attacks is the strongest single bond carbon forms, and its low polarisability and poor leaving-group ability make it resistant to the nucleophilic substitution chemistry that dehalogenases use on other halogenated compounds, e.g. chloro- and bromo-substrates ^9,10^. Chan et al. established the structural and kinetic dissection of the FAcD reaction coordinate mechanism in detail ^11^, and comparative work by Holmquist, M. ^12^ across microbial homologues showed that the characterised enzymes share an α/β-hydrolase fold, form homodimers with one substrate-access tunnel per protomer, and converge on an aspartate-histidine-aspartate catalytic triad, together with conserved residues that coordinate the substrate carboxylate and stabilise the departing fluoride ^13^. Characterised activity spans phylogenetically diverse bacteria, including *Delftia acidovorans* ^14^, the purple non-sulfur phototroph *Rhodopseudomonas palustris* ^11^, *Dechloromonas aromatica*, *Polaromonas sp. and the cyanobacterium Nostoc* sp. ^13^. Their natural repertoire, however, is narrow: of eight purified dehalogenases with confirmed FA activity, only three defluorinated difluoroacetate appreciably and none acted on perfluorooctanoic acid (PFOA) ^13^, so the enzymology that makes FAcD attractive for remediation has been demonstrated almost entirely on a single monofluorinated substrate.

In FAcD, access to the buried active site is conformationally gated ^15^, and defluorination occurs in two steps. First, the reaction proceeds through a covalent aspartyl-ester intermediate. A water molecule, activated by a histidine base, subsequently hydrolyses this intermediate, regenerating the catalytic aspartate and allowing the enzyme to enter another reaction cycle ^11,16^, with the rate-determining step being a backside S_-_2 displacement of fluoride. Hybrid quantum-mechanics/molecular-mechanics treatments characterise this as a substantial but surmountable activation barrier that rises with increased fluorination around the scissile carbon ^11,17^. Second, an S_-_2 displacement is geometrically strict: the nucleophile must approach the scissile carbon along the axis opposite the leaving fluoride, and the fraction of time a solvated enzyme spends in such a near-attack conformation influences the reaction rate ^18–21^. A *gem*-difluoro centre and a trifluoromethyl group present a sterically congested and electron-poor target in which that backside trajectory is shielded, an effect sometimes described as a Teflon shield. Hence, the substrates of greatest environmental concern are the least suitable for the native active site ^17,22^.

Together, these features suggest that screening enzymes for defluorination competence should not rest on a single static structure; it must reach the solvated ensemble and finally the quantum-mechanical (QM) cleavage step.

Protein engineering can push FAcD beyond its natural repertoire. A gate-based synergistic chain design remodelled the substrate-access channel of RPA1163 through an allosteric residue pair. It produced variants with turnover numbers above 2 × 10^5^ on the non-natural α-fluoro substrate α-fluorophenylpropionic acid ^23^, and growth-based selection has extended dehalogenase activity to additional non-natural organofluorides ^24^, together showing that the enzyme is engineerable toward other fluorinated substrates. Such efforts start from a single well-characterised enzyme and a defined mutational logic. They do not address the prior and complementary question of whether natural sequence diversity already contains variants whose active-site geometry is already suited to polyfluorinated targets and would therefore provide better starting points for engineering.

Answering that question means searching a sequence space far too large to interrogate experimentally, using a criterion that reflects catalytic competence, not binding alone. Deep-learning protein-ligand co-folding, exemplified by AlphaFold 3 ^25^ and the openly licensed Boltz models ^26^, now predicts three-dimensional enzyme-substrate complexes directly from sequence and ligand chemistry at a scale that makes library-wide screening conceivable, and related deep-learning tools for enzyme discovery and ligand-aware binding-site prediction are proliferating ^27,28^. But co-folding confidence is not the same as physical correctness; these models can place a ligand in a pocket with high predicted accuracy while failing to learn the underlying physics of the interaction, and they are biased toward their training data and insensitive to binding-site perturbations ^29^. For a catalytic question, this gap is decisive, because a well-formed but mechanistically inert pose is worthless for defluorination. Here, we use natural FAcD sequence diversity to build a mechanism-based shortlist of candidates for more highly fluorinated substrates.

In this study, we built **DeFluorX**, a single bash-initialised, end-to-end PFAS defluorinating enzyme-mining pipeline that couples deep-learning prediction of enzyme-substrate complexes to an explicit, physics-based interrogation of catalytic geometry. Using Boltz-2 ^26^, whose open weights and direct simplified molecular-input line-entry system (SMILES) input allowed a screen of this size to be run locally under a fixed seed, we modelled 2,150 bacterial FAcD variants against 27 fluorinated substrates and, rather than ranking by confidence, imposed a mechanistic filter keyed to the reactive centre: the D110 Oδ···Cα distance and Oδ-Cα-F attack angle, the catalytic triad relay and the halide cradle, penalised by the scissile C-F bond dissociation energy, backside occlusion and β-fluorination. We then subjected the top-ranked complexes to explicit-solvent molecular dynamics (MD) Simulations, endpoint Molecular Mechanics with Generalised Born and Surface Area (MM-GBSA) binding free energies, WaterMap hydration analysis, and, finally, a Quantum Mechanics/Molecular Mechanics (QM/MM) relaxed scan of the C-F cleavage coordinate. However, we intentionally did not predefine enzyme-substrate pairings, allowing candidates to be ranked by mechanistic merit rather than prior tractability assumptions.

## 2. Methods

The **DeFluorX** pipeline comprises seven major steps (Figure 1a-e): ligand and sequence curation; AI co-folding and mechanism-first tiering; statistical validation; phylogeny; structure preparation; hydration and MD Simulations with endpoint free-energy analysis; and QM/MM defluorination scanning.

**Figure 1.**
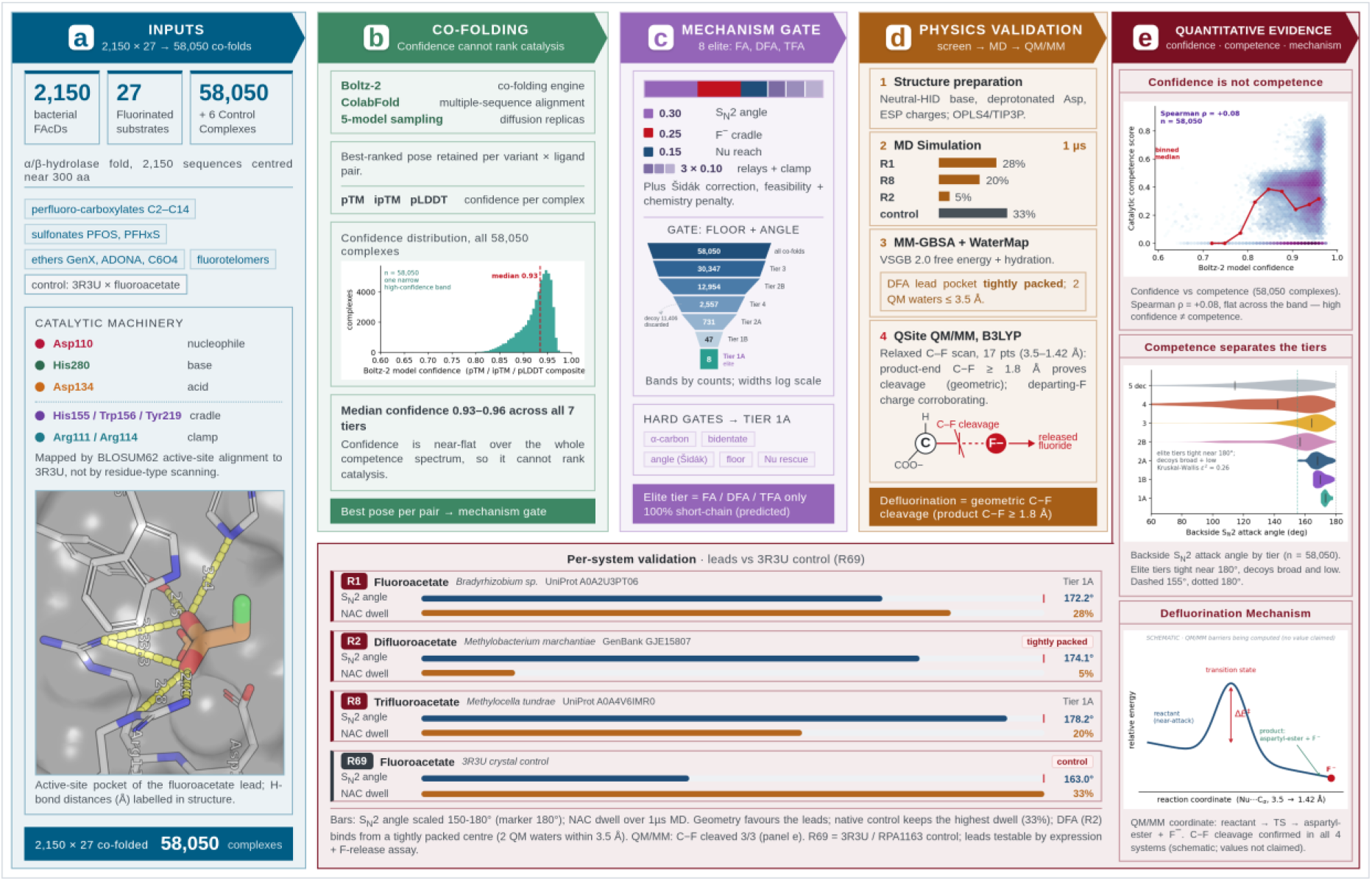
Overview of the DeFluorX pipeline. All 2,150 bacterial fluoroacetate-dehalogenase variants were co-folded against 27 fluorinated substrates with Boltz-2, giving 58,050 complexes, and filtered by a seven-tiered backside-S_N_2 mechanistic-competence gate, not by model confidence, then validated by statistical analysis, hydration-site thermodynamics, explicit-solvent MD simulations and B3LYP QM/MM scans in which defluorination is assessed by geometric C-F cleavage. **(a)** Inputs and catalytic machinery (control RPA1163 numbering). **(b)** Co-folding and model confidence. **(c)** Mechanistic-competence gate, hard gates and the tiering. **(d)** Physics validation (MD near-attack dwell, MM-GBSA and WaterMap, QSite QM/MM). **(e)** Quantitative evidence with per-system validation: per-system backside-S_N2 validation of the tier-1A leads (FA, DFA and TFA) against the RPA1163 reference, together with screen-wide evidence that model confidence does not predict catalytic competence (Spearman ρ = +0.08 with the competence score), that competence separates the tiers, and the defluorination mechanism.

### 2.1 PFAS and Protein-Sequence Curation

We encoded 27 environmentally prioritised fluorinated substrates as canonical SMILES: fluoroacetate, which is not a PFAS under the OECD definition, and 26 PFAS comprising perfluorocarboxylates and sulfonates of varying chain length, fluorotelomer alcohols and ether replacements (GenX, ADONA, C6O4) ^1,30,31^. Each substrate was encoded in the protonation state in which it was curated, not standardised to a common state: 17 as neutral carboxylic acids and 6 as neutral sulfonic acids, GenX and difluoroacetate as deprotonated carboxylates, and the two fluorotelomer alcohols as neutral alcohols. Further, the FAcD variant library was curated from bacterial sequences (13 January 2026 from UniProt and NCBI, NCBI taxonomy ID 2) ^32,33^, deduplicated by exact sequence identity and length-filtered to 250–360 aa (retrieved sequences only), reducing 15 labelled reference sequences plus 3,275 retrieved sequences to 2,150 unique variants (SS_01) that were co-folded against all 27 substrates, giving 58,050 complexes. We also defined six initial control complexes: RPA1163 (PDB 3R3U ^8^) and DeHa4 from *Delftia acidovorans*, each against fluoro-, difluoro- and trifluoroacetate to calibrate the mechanistic score against a known defluorinating system ^11,14^; for the physics-based computations, only 3R3U-FA was selected for the control.

### 2.2 Protein-Ligand Co-folding and Structural Scoring

Each FAcD variant-ligand pair was co-folded with Boltz-2 (v2.2.1, https://github.com/jwohlwend/boltz) ^26^ with the protein supplied as its amino acid sequence and the ligand as canonical SMILES, and with no templates, pocket restraints or covalent-link directives, so that every pose was generated *de novo*. We obtained multiple-sequence alignments (MSAs) for each unique protein sequence from the ColabFold MSA server (https://api.colabfold.com) ^34^. We concatenated the UniRef and BFD/MGnify/MetaEuk/SMAG alignments into a single alignment per sequence and reused it across all 27 ligand co-foldings; when the alignment server failed, we generated the alignment at prediction time instead. The six control complexes and the 3R3U ^8^ reference, together with any screen job whose batch failed and was retried individually, had their alignments resolved at prediction time, not taken from the cached alignment. Predictions were generated in two modes. Screening ran in batches of up to 20 proteins, with one recycling step and five diffusion samples per pair, using a fixed seed of 42 and writing mmCIF output with the custom CUDA kernels disabled ^26,34^. The five diffusion samples per pair are independent samples from the single Boltz-2 model, not an ensemble of five models. We selected the representative pose across the five diffusion samples tier-first: the highest degrader tier reached by any sample, then the largest raw S_-_2 attack angle (rounded to 0.1°), then the highest competence score, with Boltz-2 confidence as the final tie-break. Reported geometries are therefore best-of-five selections on the same criteria as the ranking scores. The fraction of the five samples that independently reached a degrader tier was recorded as a per-complex consensus value and used as the final tie-break in the mechanism-first ranking, after tier, competence score, catalytic assembly score and active-site conservation, so a pose found by one sample ranks below an otherwise equal pose found by several without ever crossing a tier or competence boundary. We retained all five samples for variance analysis, and we tabulate a subset in SS_06. A composite interaction score was computed for each retained complex:

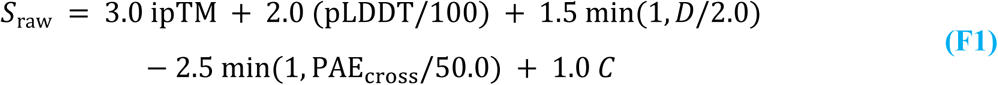

where *D* is the interaction density (the number of detected interactions divided by the number of ligand heavy atoms, saturating at 2.0) counted by the pipeline’s geometry engine using criteria that reproduce the Schrödinger Maestro (https://www.schrodinger.com/) default interaction definitions ^35–45^, and PAE_cross_is the mean cross-interface predicted aligned error, and *C* is the Boltz-2 ^26^ global confidence score ^25^. The raw sum of F1 ranges from −2.5 to +7.5, is centred at 2.5, is divided by a gain of 1.25, and is transformed into a bounded composite interaction score by:

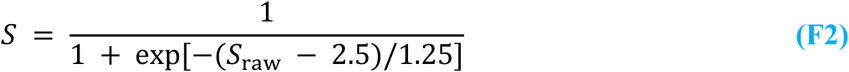

The logit is clamped to ±50 before the sigmoid, a guard that does not bind for any complex in this screen. The five weights, the centring and the gain were assigned a priori to rank the confidence signals by expected reliability; they were not fitted to experimental binding data, so *S* is a bounded ranking score, not a binding probability. Four of the five terms (ipTM, pLDDT, the global confidence and the cross-interface predicted aligned error) are Boltz-2 confidence outputs, and interaction density is the only term computed from the pose itself, carrying 1.5 of the 7.5 total positive weight. The score is therefore a confidence-dominated composite by construction, and it is used only as a Pareto objective alongside the mechanistic score, never as evidence of reactivity. The weights were set a priori rather than fitted because no labelled binding or activity data exist for this enzyme-substrate space at this scale; fitting them would require a target, and the only abundant target available is the co-folding confidence the analysis sets out to interrogate, so a fitted score would encode the very signal it is meant to be tested against.

### 2.3 Mechanism-First Ranking Hierarchy

Catalytic machinery was mapped onto every complex by BLOSUM62 pairwise alignment (*Bio*. *Align*. *PairwiseAligner*, *open* − *gap* − 10.0, *gap* − *extension* − 0.5) ^46^ against the PDB 3R3U reference numbering ^11^, never by residue-type scanning ^47^. All eight active-site positions were resolved this way: the aspartate nucleophile D110, the arginine carboxylate clamp R111 and R114, the aspartate acid catalyst D134, the halide stabiliser H155, the halide cradle W156 and Y219, and the histidine base H280 ^8,11,13^. We flagged a mapping as reliable when the pairwise alignment reached at least 25% sequence identity. We also banded each alignment by sequence identity into nine grades, from A (above 90%) to I (at or below 20%), in ten-point intervals that are left-open and right-closed; the grade accompanies every complex in the accompanying tables. Where the nucleophile fell just outside its aligned column, a rescue rule recovered it, but only within two residues for a complex to remain eligible for the elite tier, so that a distant aspartate cannot stand in for the true nucleophile. We list complexes whose active site could not be mapped in SS_03.

A configuration-weighted mechanistic score combined the nucleophile reach (D110 Oδ to substrate α-carbon, weight 0.15) ^19–21^, the Nuc-Base and Base-Acid relay distances (0.10 each, gated at 5.0 and 5.5 Å), the bidentate carboxylate clamp (0.10), halide stabilisation (0.25) and a graded S_-_2 attack angle (0.30) ^18,48^, the angle entering as the Šidák-corrected effective angle of F5, not the raw angle ^49^. The correction is applied once, before scoring: the raw angle and the fluorine count on the scissile carbon are converted to an effective angle, which both the angle threshold of the tier scheme and the graded angle term then read, so a polyfluorinated substrate is not penalised twice for the same property. The exponent in F5 is the fluorine count on the scissile carbon; there is no factor of two for the two aspartate oxygens, because the joint near-attack rule fixes the attacking oxygen. Steric clashes were penalised by 0.03 points each, to a maximum of 0.15. A parallel competence score weighted attack angle 0.30, nucleophile distance 0.25, clamp 0.20, trajectory 0.10, triad 0.10 and halide contact 0.05. A third, fully continuous soft catalytic score was computed alongside these two, replacing every binary anchor with a logistic so that a complex is not moved across a boundary by a hair’s-breadth threshold crossing; it gates nothing and ranks nothing, and is reported as a fully continuous companion to the mechanistic score and as the quantity underlying the catalytic-geometry figures, with its definition given as SF18 in the Supplementary Methods (SF1–SF18). Reactive-centre gating required attack on the α-carbon adjacent to the carboxylate. Graded chemistry and containment penalties were then subtracted from the geometric score to give the effective score on which the tier thresholds apply: 0.015 per kcal mol⁻¹ of scissile C-F bond dissociation energy above 123.0 kcal mol⁻¹, 0.13 per Å of backside occlusion above a 2.0 Å van der Waals sum ^50^, and 0.08 per unit of β-withdrawal on the attack-carbon environment, where each vicinal fluorine and each ether oxygen counts as one unit. Only the occlusion term fades as the trajectory approaches linearity, from its full value at or below 175° to zero at 180°, on the reasoning that a pose reaching the anti-periplanar axis has already cleared the obstruction that term measures; bond strength and β-withdrawal are properties of the substrate and keep their full weight at any angle. Where the scissile centre was unresolved, or the backside occlusion could not be measured, the corresponding term was set to zero, not imputed, and the complex was flagged and barred from the elite tier, so that a mapping failure could not invent a penalty; all 405 complexes whose nucleophile could not be resolved were classified as decoys. Chemical feasibility does not enter this score at all: it is a multiplicative factor on the competence score (F6, F12), floored at 0.10, so a recalcitrant substrate is held near, never below, the bottom of the feasibility range and is ranked on merit, not removed by a substrate-class veto ^9,10^.

Complexes were then placed on seven tiers (Tier_1A, Tier_1B, Tier_2A, Tier_2B, Tier_3, Tier_4, Tier_5_Decoy) by explicit thresholds, not by clustering. The tier names label positions in a geometric scoring hierarchy, not claims of measured activity; “degrader tier” throughout means a complex that satisfies the mechanistic criteria for Tiers Tier_1A to Tier_2B. Tier_1A required a nucleophile distance ≤3.0 Å, an effective attack angle ≥170° after the Šidák correction of F5, a Nuc-Base distance ≤3.5 Å, a Base-Acid distance ≤4.5 Å, an effective mechanistic score ≥0.85, a scissile C-F bond dissociation energy ≤128.0 kcal mol⁻¹, and an active-site predicted local distance difference test (pLDDT) score ≥90 or, where that was unavailable, a global Boltz-2 ^26^ confidence ≥0.85. A pose meeting these thresholds held Tier_1A only if it also cleared a downgrade-only machinery gate: either complete catalytic machinery with an effective mechanistic score ≥0.90, or near-complete machinery with an effective mechanistic score ≥0.85 redeemed by a catalytic assembly score ≥0.74 (F13), so that a slightly backside-occluded substrate can register as elite only in its single most crystal-exact pose. SS_02 provides full per-tier thresholds and every variant sequence, its source accession, assigned tier, Boltz-2 metrics and derived scores.

### 2.4 Pareto Optimisation

Pareto non-dominated sorting was run on two objectives, the Boltz-2 ^26^ global confidence score and the composite interaction score of F2, over catalytically viable poses only, defined as an effective mechanistic score at or above the Tier_2A floor of 0.70; non-viable poses were held off the front, not ranked on it, so that no inhibitor-like pose could present as optimal (SS_04). Because F2 embeds the same confidence signals, this describes the trade-off within the confidence-derived scores, not an independent test of confidence against reactivity, and hierarchical clustering of alignment-free k-mer cosine distances (k = 3, unweighted pair group method with arithmetic mean linkage) was used to visualise how the tiers distribute across the sequence library (SS_07); the dendrogram is a sequence-similarity tree and carries no substitution model. Principal-component analysis was run on eight descriptors, the Boltz model confidence, ipTM, mean pLDDT, normalised interaction density, chemical affinity, binding probability, hydrogen-bond count and salt-bridge count, and is reported as per-feature loadings on the first two components (SS_04); the loadings are unit-normalised eigenvector components and therefore carry direction only, so no explained-variance fraction is reported. Chemical-space structure was visualised as a uniform manifold approximation and projection (UMAP) embedding computed with n_neighbors = min(15, n − 1), min_dist = 0.1 and a fixed random state of 42, displayed as a hexbin coloured by mean competence with the elite tier overlaid;

UMAP distances are not metric, so the embedding supports statements about which substrates cluster together and none about the magnitude of the separations ^51^. We also cross-classified complexes by tier against Boltz-2 confidence to separate confidence-reactivity conflicts (SS_04). Among degrader-tier complexes (Tier_1A to Tier_2B), those with confidence at or above 0.70, a cut-off chosen for this classification, are competent-and-confident, which the classification code labels Consensus High, and those below it are competent low-confidence, which the classification code labels Hidden Gem. Among Tier_4 and Tier_5_Decoy complexes, those at or above 0.70 are Decoy, and those below 0.60 are incompetent-and-doubted, which the code labels Consensus Low, and the cut-offs differ, with 0: Every other complex, including all of Tier_3, stays in the ambiguous class, which is where a complex ends up when no rule assigns it rather than the outcome of a test. Cross-tier hypothesis tests used the Kruskal-Wallis test with pairwise Mann-Whitney statistics ^52,53^, and Benjamini-Hochberg false-discovery-rate control ^54^ was applied at the end of the run across a single family of 96 tests, comprising the cross-tier and degrader versus non-degrader comparisons, a chi-square test of interaction-density band against tier, a Wilcoxon signed-rank paired test, a silhouette label-permutation test of tier separation in the UMAP embedding over 199 permutations with the sample capped at 2,000 complexes, the remaining panel and battery tests and the pairwise correlations among the six ranking metrics, all corrected in one pool and reported in SS_05. With 199 permutations, the smallest p-value this test can return is 1/200 = 0.005. The other tests in this family use asymptotic approximations and return much smaller p-values, some below the limit of double precision and recorded as zero. The permutation result therefore cannot be compared by p-value, and its Benjamini–Hochberg q-value cannot be below 0.005. We corrected the feature-correlation matrix separately using only its unique upper-triangular pairs so that the same Spearman statistic can have different q-values across the two analyses. To avoid inflating the correction, we excluded underpowered tests upfront: degrader comparisons required ≥30 observations, and feature pairs required ≥20 jointly observed complexes. The two sample-size floors above and the within-run tercile cutoff used to classify physics-strong and AI-strong complexes are not constants and are listed in Supplementary Table S1. Because the per-tier distributions of the geometric descriptors are non-normal, we used non-parametric comparisons throughout. The Kruskal-Wallis *ε*^2^ effect size (F3) quantified magnitude, which expresses the fraction of rank variance explained by the tier factor, with *k* the number of tiers and *n* the total number of complexes:

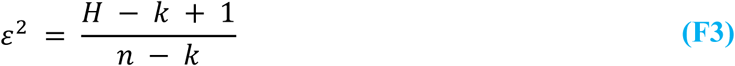

Because the tiers are defined by thresholds on these same descriptors, the cross-tier comparison is descriptive rather than a test of separation; the effect sizes are reported to quantify how much of the rank variance the tier factor accounts for: the effective mechanistic score differs across tiers with *ε*^2^ = 0.47 and the raw mechanistic score with *ε*^2^ = 0.40, the S_-_2 attack angle with *ε*^2^ = 0.26 and the nucleophile distance with *ε*^2^ = 0.11 (Kruskal-Wallis ^53^ *p* underflowed to zero in double precision, below approximately 5 × 10^−3^^24^ , in every case, *k* = 7, *n* = 58,056 complexes including the six calibration controls), so the hierarchy reflects a substantial fraction of the total variance in reactive geometry, not a marginal, sample-size-inflated significance. The same Šidák multiplicity correction used for the attack angle (F5) controls the family-wise error introduced by the multiple equivalent C-F bonds of a polyfluorinated carbon. Numbers-of-sequences, tier counts and geometric medians are taken directly from the ranked complex-level table, and the full statistical output is provided in SS_05.

We extracted source organisms and their taxonomy from the sequence accessions: the fluoroacetate lead from *Bradyrhizobium* sp. (UniProt A0A2U3PT06), the difluoroacetate lead from *Methylobacterium marchantiae* (GenBank GJE15807), and the trifluoroacetate lead from *Methylocella tundrae* (UniProt A0A4V6IMR0), all Alphaproteobacteria.

### 2.5 Scoring Formulae

The mechanistic score is a configuration-weighted sum of six reactive-geometry components with weights summing to one (F4, Figure 1c). Five are binary anchor checks, each contributing its full weight when it meets its threshold. These are the nucleophile near-attack reach *g*_nuc_ (Oδ-Cα ≤3.2 Å), the nucleophile-base and base-acid triad relays *g*_NB_ and *g*_BA_ (≤5.0 and ≤5.5 Å), the bidentate carboxylate clamp *g*_clamp_ , and the halide stabilisation *g*_stab_ . The residue identities and catalytic roles behind these five were established for fluoroacetate dehalogenase by crystallography and mutagenesis ^8^ and by reaction-coordinate mapping ^11^, and the reach threshold follows the near-attack-conformation criteria of Lightstone and Bruice ^19^. The backside attack angle *g*_angle_ , is the Šidák survival evaluated at the effective angle of F5, *g*_angle_ = [1 + cos(180° − *θ*_eff_)] / 2, which is identically the survival term a of F12; its angular dependence follows the nucleophilic-approach trajectory of Bürgi, Dunitz and Shefter ^18^ and Bürgi et al. ^48^. The six components combine as:

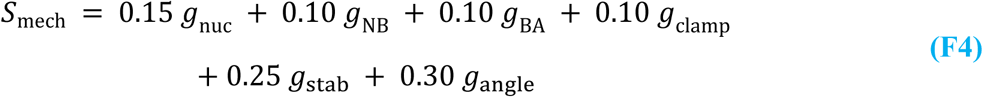

The component definitions, the interaction-affinity and WaterMap-water formulae, and all weights and thresholds are provided in full in the Supplementary methods (SF1-SF18 and Supplementary Table S1). The angle term is evaluated on a multiplicity-corrected attack angle so that a polyfluorinated carbon presenting n equivalent scissile C-F bonds is not rewarded for the single most favourable of n independent chances. Writing the angular defect from the ideal 180° backside geometry as a per-bond miss probability p1, the n bonds are combined by a Šidák survival correction ^49^ and back-transformed to an effective angle (F5): The correction is not a claim that several bonds react: the reaction proceeds at one C-F bond, and the correction applies to the selection statistic, because taking the maximum angle over n equivalent bonds is a biased estimator of the alignment of the bond that reacts.

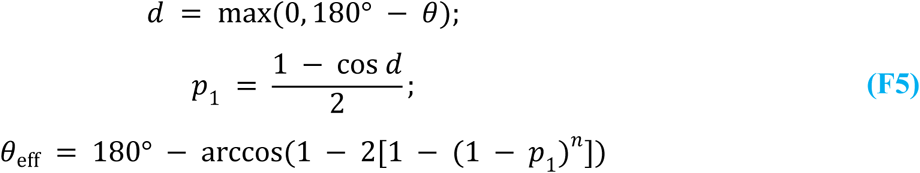

Chemical feasibility enters as a continuous, never-vetoing factor (F6). The leaving-group term *f*_*BDE* falls linearly from one to zero as the scissile C-F bond-dissociation energy rises across the 120 to 132 kcal mol⁻¹ window, following the established dependence of backside S_-_2 leaving-group ability on C-F bond strength ^9,10^. The β-withdrawal term *f* discounts vicinal fluorination and ether oxygens (*n_β_*), which withdraw electron density from the reactive centre ^9^ and track the substrate range of the characterised enzymes, which defluorinate difluoroacetate but not perfluorooctanoic acid ^13,17^. Their product is floored at 0.10, so a recalcitrant substrate is scaled down in rank but never excluded. The two terms are defined as:

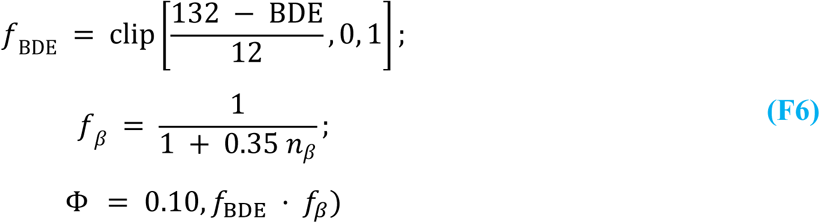

The tier thresholds and the elite gate use an effective score that subtracts the graded chemistry and containment penalties from the geometric score and is floored at zero (F7, F8 and F11). *O*_bs_ is the backside steric occlusion in Å, and s is a linear fade, full at or below 175° and zero at 180°, that applies to the occlusion term alone. Occlusion obstructs the backside trajectory itself ^10^, so a straightening approach relieves it. The bond-strength and β-withdrawal terms do not fade, because neither is an obstruction that a favourable trajectory can relieve:

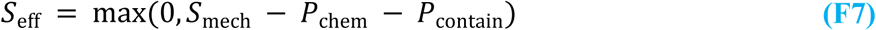

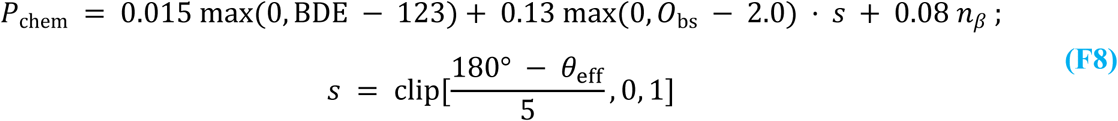

A trajectory frame is scored as a near-attack conformation when the catalytic triad is intact, and the aspartate-oxygen to α-carbon distance and the backside attack angle simultaneously satisfy the strict gates (Figure 1d, Desmond MD). The distance and angle criteria follow the near-attack-conformation formalism ^19^. Occupancy is the fraction of post-equilibration frames that qualify; frames recorded before the box equilibrated are flagged as such in the per-frame record and excluded from both the numerator and the denominator. The occupancy is distinct from the longest continuous residence reported in Table 3 (F9):

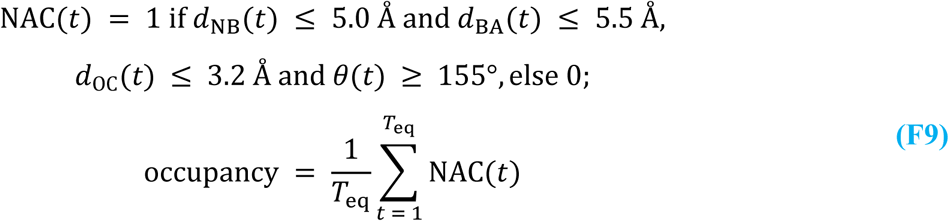

The per-hit turnover call is the geometric criterion for C-F scission (Figure 1d, QSite QM/MM). The scissile bond must measurably break in the QM/MM scan, with its product-end length reaching 1.8 Å, a geometric criterion defined in this work at approximately 1.3 times the reactant-state C-F bond length and corroborated by the departing-fluoride charge (F10). The departing-fluoride Mulliken charge ^55^, which under-reports the free-ion value in the diffuse-function-free basis, and the one-dimensional-scan barrier are reported as corroborating context, not as independent conditions:

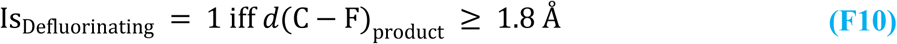

The containment penalty of F7 charges a complex whose cavity fails to enclose the ligand. It takes the cavity-containment fraction *C*_cav_ and penalises any shortfall below 0.85 at unit weight (F11):

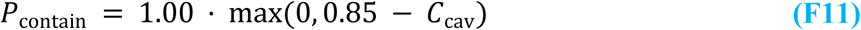

The parallel competence score named in section 2.3 gates Tier_2A and Tier_2B, with floors of 0.50 and 0.40; the two elite tiers gate on geometry, on the machinery gate and on the catalytic assembly score, and do not read it. The same two floors are reapplied after those caps, so a pose demoted into Tier_2A or Tier_2B rather than entering it on the forward path drops to Tier_3 and loses degrader status if its competence sits below the floor. The score is given in full here, and its six terms carry different weightings and a different composition from F4. The nucleophile distance rises from 0.15 to 0.25 and the clamp from 0.10 to 0.20, halide stabilisation falls from 0.25 to 0.05, and the attack angle is unchanged at 0.30. The two separate proton-relay terms of F4, 0.10 each, are merged into a single triad term that carries 0.10 in place of their combined 0.20 and is evaluated as the mean of the two graded relay distances; a trajectory term is added, for which F4 has no counterpart. Both scores have six terms because the merger makes room for the trajectory term. The feasibility factor of F6 scales the weighted sum, and the angle term here is the Šidák survival itself rather than the back-transformed effective angle of F5 (F12):

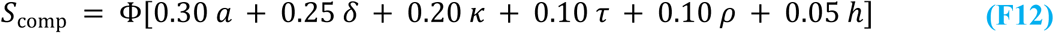

The six components are defined in the Supplementary Methods (SF14): *a* the Šidák survival, *δ* the nucleophile distance, *κ* the clamp integrity, *τ* the trajectory deviation, *ρ* the triad relay and ℎ the halide-stabilisation indicator. The symbol *δ* here is the nucleophile-distance term and is distinct from *d*, the angular defect of F5.

A catalytic assembly score caps the tier a complex can attain. It measures how closely the eight active-site positions superpose on the 3R3U reference ^11^, as the reciprocal of one plus the active-site root-mean-square deviation (RMSD), so an exactly superposed active site scores one (F13). The component definitions are given in the Supplementary Methods:

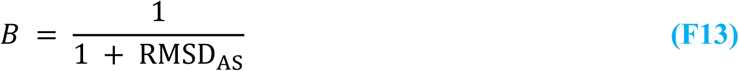

### 2.6 Structural Refinement and Protonation-State Assignment at pH 8.0

The MD cohort was selected as the best complex per unique ligand, drawn only from Tier_1A, and flagged in the ranked table (SS_02). Only that cohort was carried forward to all-atom preparation with Schrödinger PrepWizard at pH 8.0, within the alkaline range in which the reference enzyme RPA1163 is characterised and assayed ^11^, using PROPKA protonation and Epik ligand protonation at the same pH ^56^, and the S-OPLS force field, with clash-resolving minimisation restrained to 0.15 Å RMSD ^57,58^. The catalytic aspartates were then overridden to the deprotonated state by removing the carboxyl proton, and the histidine base to the neutral HID tautomer (*Nδ*1 − *H* present, *Nε*2 free) required for its general-base role, irrespective of the predicted pKa; this enforcement is material, because the protonated HIP form collapses the S_-_2 attack angle during relaxation. Because the aspartate protonation is set explicitly in this way, the mechanistic conclusions do not depend on the precise pH value. Backbone *φ*/*ψ* distributions were computed before and after preparation, and for the predicted models against the reference crystal structure, as a check that protonation-state assignment and restrained minimisation left the fold geometry unperturbed. We derived electrostatic-potential-fitted charges for the reactive ligands as B3LYP/6-31G** single points in Jaguar to parameterise the subsequent simulations ^59–61^. Pose geometry before and after preparation, including the backside attack angle in each state and the change between them, is tabulated in SS_08, and the fitted charges are listed in SS_09; the prepared poses and their interaction maps are shown in Supplementary Figure 5.

### 2.7 Molecular Dynamics, Hydration Thermodynamics and Endpoint Binding Free Energies

All prepared the 3 Tier_1A identified unique best complexes with control were further solvated in an orthorhombic three-site transferable intermolecular potential (TIP3P) box, neutralised with counterions and 0.15 M NaCl, with ion placement excluded within the reactive pocket to avoid screening the nucleophile, and simulated with Desmond under the OPLS4 force field in the isothermal-isobaric (NPT) ensemble at 300 K ^62–64^. Ligand heavy atoms were positionally restrained at 5.0 kcal mol⁻¹ Å⁻² and the protein backbone at 2.0 kcal mol⁻¹ Å⁻² throughout production, so pocket retention is restraint-enforced, not spontaneous. The restraint was applied because the reactive pose is the object under test: in unrestrained trials, a small substrate such as fluoroacetate relaxed out of the near-attack geometry and diffused into bulk within the first nanoseconds, after which every downstream read-out correctly reported an empty active site. A low, flat ligand RMSD therefore indicates that the restraint held rather than that the pose is intrinsically stable. Every near-attack occupancy reported in this work is therefore restraint-conditioned. We analysed production trajectories for catalytic-triad and halide-cradle stability and for near-attack-conformation occupancy, applying the near-attack criteria to the mapped aspartate oxygen, the *Cα* and the backside-attack trajectory ^19^. All interatomic distances measured on the trajectory were computed under the minimum-image convention with respect to the periodic box. Each selected system was run as a single one-microsecond (1,000 ns) production trajectory (n = 1 replicate) under the OPLS4 and TIP3P setup; build and solvation quality control for these systems is summarised in Figure 6a. The production run was kept under the Martyna-Tobias-Klein method ^65^, with thermostat and barostat relaxation times of 1 and 2 ps and isotropic pressure coupling to 1.01325 bar, on RESPA ^66^ timesteps of 2 fs for the bonded and near-field terms and 6 fs for the far-field term, writing at every 10 ps totalling 100,000 frames, with velocities randomised once from a fixed seed of 2024. Equilibration was detected, not assumed: the trajectory was divided into 200 blocks, the first fifth discarded, and a block was accepted as equilibrated when the temperature sat within 3 K of the 300 K target, the box volume within 1% of its mean with a drift below 0.05% per nanosecond, and the *Cα* RMSD below 3 Å; the equilibrated window begins at the first point from which at least 90% of subsequent blocks pass. All reported near-attack occupancies are computed over that window only. Per-frame near-attack data for every trajectory are tabulated in SS_13.

WaterMap grand-canonical hydration analysis, run for 5 ns, identified the hydration sites of the reactive pocket and their thermodynamic signatures. The ligand was retained so that the analysis describes the occupied holo pocket relevant to catalysis, not apo displacement, and the site free energies were interpreted alongside the binding energetics ^67^. Site-level occupancies, enthalpies, entropies, free energies and hydrogen-bond counts are tabulated in SS_10.

Prime MM-GBSA endpoint binding free energies were computed with the variable-dielectric surface generalised Born (VSGB) 2.0 implicit-solvation model and aggregated per rank from the MD ensembles ^68,69^. Sampling covered a 100 ns window of each trajectory at every tenth frame, averaged as the mean, with 95% confidence intervals from a block bootstrap of 2,000 resamples using 50-frame blocks. Per-rank aggregates, including Boltzmann-weighted mean, median, minimum, maximum and standard-deviation values, are tabulated in SS_11.

### 2.8 Quantum-Mechanical Transition-State Analysis and NAC Estimation

Relaxed scans drove the forming nucleophile-oxygen to substrate-carbon coordinate from 3.5 to 1.42 Å in 17 steps; the scissile C-F bond was not itself a scan variable, so its elongation is observed as a consequence of the driven nucleophilic approach, not from an independent transition-state search. A product-end scissile C-F length at or beyond 1.8 Å registers cleavage, the basis-robust criterion, with the departing-fluoride Mulliken charge (which under-reports the free-ion value in the diffuse-function-free 6-31G** basis, uncorrected for dispersion) retained as a corroborating read-out ^55^. The QM-region net charge was derived per system by summing the formal charges the prepared structure assigns to the QM atoms, rather than assuming expected protonation states; for the carboxylate substrates in this cohort, it evaluates to -3, from the two deprotonated catalytic aspartates and the substrate carboxylate, with the histidine base neutral. The frozen-orbital treatment then removes one electron per *Cα* − *Cβ* cut, so the charge entering the self-consistent field calculation is lower than the formal charge by the number of cuts. The one-dimensional-scan energy maximum on the reactant side is reported as a barrier estimate but is not transition-state-verified and is not physically interpretable below about one kcal mol⁻¹ ^70^; a frame whose energy maximum fell outside the scan window resolved no transition state and is excluded before the barrier and reaction-energy aggregates are formed, so that a system may contribute one frame or three; because the three scanned frames are pre-selected before any scan was run in two stages: a pre-organisation score (100.0 per Å below the relaxed near-attack distance and 5.0 per degree above the relaxed angle, less 50.0 per water-blockade unit and 10.0 per kcal mol⁻¹ of unfavourable WaterMap ΔG) acts as an eligibility mask, admitting only post-equilibration frames that score above zero and in which no counter-ion approached within 3.0 Å of the nucleophile or the substrate head group, and the surviving pool is then ranked on the per-frame consensus value, with a first-shell-water preference allowed to reorder the leading candidates so that a dewetted but well-aligned frame does not win, and the control enzyme returns the highest apparent barrier, (the barrier does not rank the leads or discriminate competence), and, over converged frames only, the scan reaction energy is uphill for the fluoroacetate lead (+2.0 kcal mol⁻¹), the trifluoroacetate lead (+5.6 kcal mol⁻¹) and the reference (+8.3 kcal mol⁻¹), and downhill only for the difluoroacetate lead (−6.5 kcal mol⁻¹). The near-attack geometry conditions the reported context only, not the cleavage call. First-shell quantum waters were selected within 3.5 Å of the reaction centre and capped at three, preferring thermodynamically stable WaterMap ^67^ sites where available; every system carried two to three such waters, so the departing fluoride’s first-shell solvation is treated quantum-mechanically throughout. The calculations used QSite at B3LYP/6-31G** with an OPLS_2005 MM region and hydrogen-capped *Cα* − *Cβ* cuts; the QM region, at net charge -3, comprises the substrate, the aspartate nucleophile, the histidine base, the aspartate acid, the halide-stabilising residue and the halide-cradle residues, together with the QM waters within 3. 5 Å, capped at eight residues in that priority order ^71,72^. The MM region runs under

OPLS_2005 instead of the OPLS4 used for the trajectories: QSite exposes only OPLS_2005 and OPLS3e, and OPLS3e aborts on a frozen-orbital cut, so the QM/MM force field does not match the force field used to generate the ensemble, which is a limitation of QSite ^73^. The relaxed-scan maximum is therefore reported as a scan feature, not as an activation energy, and is not used to rank the leads; the established backside-S_-_2 mechanism of FAcD provides the reference reaction coordinate ^11^. We tabulate the per-rank defluorination geometry in SS_12, and the per-frame QM/MM scans from which these aggregates form in SS_14.

### 2.9 Data and Code Availability and Computational Environment

The DeFluorX pipeline code is available at https://github.com/KU-MGB/DeFluorX and is permanently archived at Zenodo: https://doi.org/10.5281/zenodo.23318610. Structure cofolding used Boltz-2 ^26^ (model boltz2) with ColabFold multiple-sequence alignments. We used Schrödinger Release 2026-1 for MD Simulations ^62^, WaterMap ^67^, Prime MM-GBSA ^68,69^, and QSite QM/MM calculations. To ensure computational reproducibility, we ran all calculations on the same workstation running Ubuntu 24.04.5 LTS, equipped with an AMD Ryzen 9 9950X3D 16-core processor and 64 GB of RAM. GPU-accelerated tasks (Co-folding, MD Simulation, WaterMap, etc.) used an NVIDIA GeForce RTX 5090 GPU (32 GB VRAM) running CUDA Toolkit 13.0. All analyses, including post-simulation analysis, used Python 3.10 with NumPy 1.26, SciPy 1.13, pandas 2.3, matplotlib 3.10 and MDAnalysis 2.9 for trajectory handling.

## 3. Results

### 3.1 A Conserved FAcD Library Enables Screening against 27 PFAS

We built the screen from a curated, deduplicated library of 2,150 bacterial fluoroacetate-dehalogenase sequences and a panel of 27 fluorinated ligands, along with six calibration controls based on the RPA1163/3R3U crystal enzyme and the functional DeHa4 homologue against fluoro-, difluoro-, and trifluoroacetate ^14^. The ligand panel spans the classes that dominate the contamination problem: short-and long-chain perfluorocarboxylates and sulfonates, fluorotelomer alcohols and the ether replacements GenX, ADONA and C6O4. Co-folding every variant-ligand pair yielded 58,050 predicted complexes. The variant library is structurally homogeneous, with a mean sequence length of 298 residues (median 296, range 194–360) and a unimodal length distribution consistent with a single conserved α/β-hydrolase architecture (Figure 2a; the variant phylogeny resolved by competence tier in Figure 2c; alignment grades, tier composition and the scope diagnostics are in Supplementary Figure 1). Catalytic-machinery mapping was near-complete: the aspartate nucleophile, the aspartate acid and the histidine base were resolved in 2,110 of the 2,150 variants (98.1%), and all eight active-site positions in 1,864 (86.7%), the tryptophan of the halide cradle being the least often resolved at 92.8% (Figure 2b; the variants missing each position are listed in SS_03). Because the scaffolds are architecturally equivalent, the differences that the pipeline ranks on arise from active-site geometry and reactive-centre chemistry, not from gross fold divergence.

**Figure 2.**
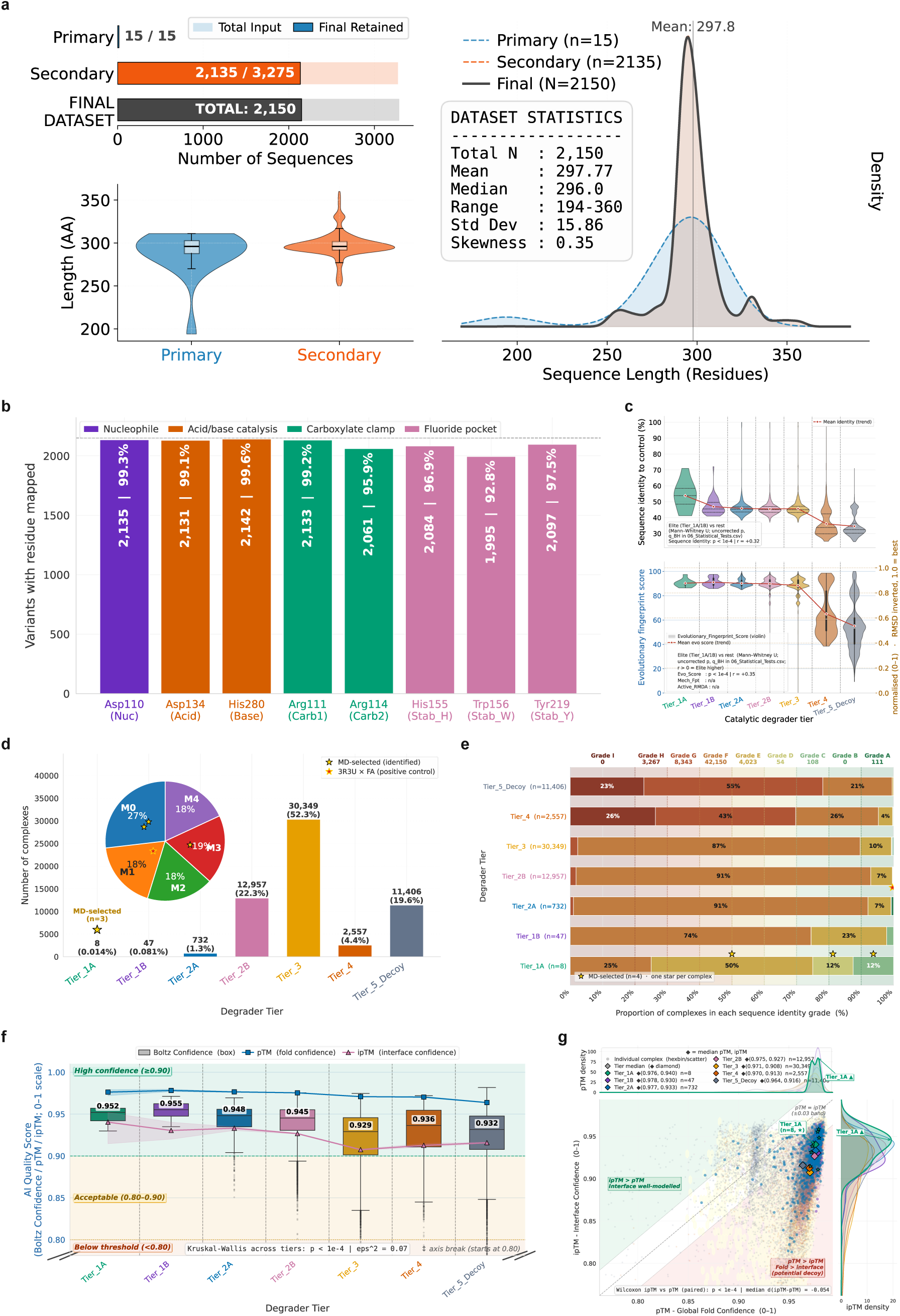
Screening and model quality. The screen and its inputs. **(a)** The curated FAcD variant library of 2,150 bacterial fluoroacetate-dehalogenase sequences co-folded against the 27-member fluorinated substrate set, with the sequence-length distribution of the primary and secondary sets. **(b)** Coverage of the catalytic-machinery mapping across the library: the fraction of the 2,150 variants to which each reference active-site residue was assigned by BLOSUM62 alignment to the 3R3U numbering (the Asp110 nucleophile, Asp134 acid, His280 base, the Arg111/Arg114 carboxylate clamp and the His155/Trp156/Tyr219 fluoride pocket), rather than by residue-type scanning. **(c)** Evolutionary phylogeny of the variant set, resolved by competence tier. **(d)** Competence-tier distribution of the 58,050 co-folded complexes, from the elite Tier_1A to the Tier_5_Decoy set; the three MD-selected leads (R1, R2, R8) and the RPA1163/3R3U control (R69) are marked. **(e)** Active-site alignment grade, A to I, relative to the 3R3U reference, resolved by tier. **(f)** Boltz-2 model confidence, pTM and ipTM by tier, showing confidence is near-flat across the full competence spectrum. **(g)** Per-complex pTM against ipTM, coloured by tier. Source data in SS_02; further dataset diagnostics in Supplementary Figure 1.

### 3.2 Boltz-2 Confidence Is High but Does Not Rank Predicted Catalytic Geometry

Boltz-2 co-folded the complexes with consistently high global model confidence, and that confidence was almost flat across the entire competence spectrum: the median Boltz confidence spanned only 0.929 to 0.955 across all seven tiers, with 0.952 in Tier_1A and 0.955 in Tier_1B against 0.936 in Tier_4 and 0.932 in Tier_5_Decoy, and the fold and interface confidences (pTM, ipTM) were similarly elevated even in the decoy set (Figure 2f, g). In other words, the model was equally confident about complexes that are catalytically poised and about complexes that are mechanistically inert.

This is the study’s central methodological observation, and it is consistent with the emerging consensus that co-folding models do not reliably learn the physics of protein-ligand interaction and are prone to confident but physically unrealistic poses ^29^. A ranking built on co-folding confidence would therefore be dominated by well-formed but reaction-incompetent complexes. The remainder of the pipeline provides the orthogonal, physics-based criterion that the AI model does not.

### 3.3 Model Confidence Does Not Discriminate between Catalytically Competent and Inert Complexes

Cross-classifying every complex on mechanistic tier against co-folding confidence resolves the dissociation quantitatively (Figure 3a). Three of the five conflict classes are populated: Ambiguous with 30,412 complexes (52.4%), Decoy with 13,898 (23.9%) and the competent- and-confident class, which the classification code labels Consensus High, with 13,740 (23.7%). The competent low-confidence class, which the classification code labels Hidden Gem, and Consensus Low are both empty. Each requires a complex whose confidence falls below a threshold of 0.70 and 0.60, respectively, and no complex in the library does: the median confidence never drops below 0.929 in any tier. Confidence produces no false negatives because it never scores a mechanistically competent complex low: not one mechanistically competent complex falls below the threshold, while 13,898 complexes (23.9% of the library) are endorsed at high confidence and classed by the mechanistic gate as Tier_4 or Tier_5_Decoy. The same asymmetry is visible directly in the confidence-against-attack-angle plot (Figure 3b, e). A confidence-ranked screen would therefore pass nearly all of the library rather than mis-order it. Consensus High is exactly the four degrader tiers, with no member outside them, and because the high-quality set is defined as those four tiers, the whole of Tier_3 falls into Ambiguous by construction, not by any test; the 65 Tier_4 and Tier_5_Decoy complexes whose confidence lies between 0.60 and 0.70 default there as well. The Decoy class is not an independent check on the mechanistic gate, since the gate itself removes those complexes, so the two classifications agree by design, not by corroborating one another. These counts come from SS_04, whose per-tier populations differ from the ranked table by a single complex at the Tier_2B/Tier_3 boundary; the totals agree at 58,056. Within the degrader tiers, the assignment is reproducible across diffusion samples: the mean fraction of the five samples independently reaching a degrader tier falls from 0.88 in Tier_1A and Tier_1B to 0.73 in Tier_2A and 0.56 in Tier_2B (Figure 3c, d, f, g; Supplementary Figure 4), so elite assignments rest on most samples rather than on one. The comparison does not extend to the non-degrader tiers, where the consensus is necessarily zero: the reported tier is the highest reached by any sample, so no sample reached a complex outside the degrader tiers.

**Figure 3.**
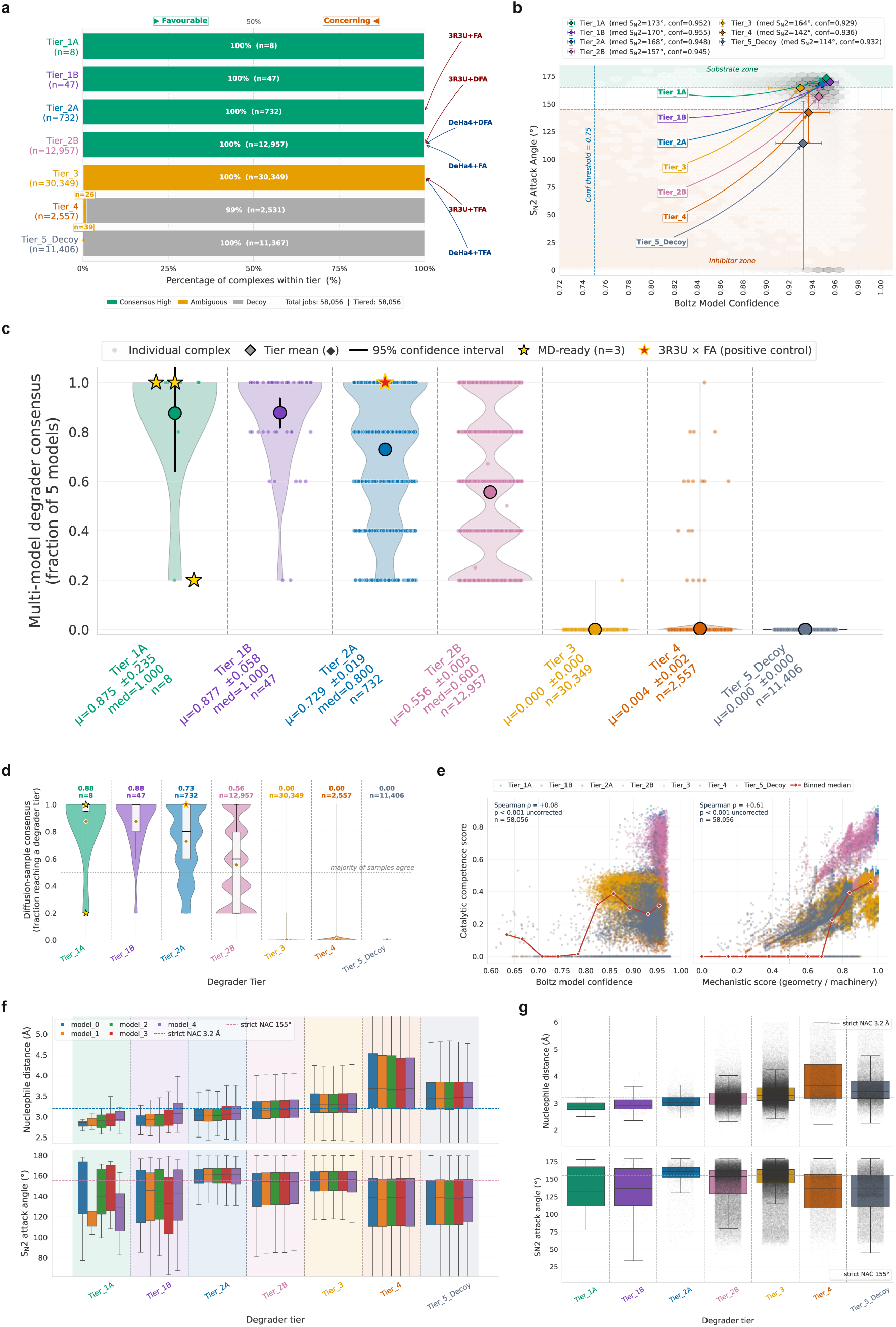
Confidence does not predict competence. The conflict between co-folding confidence and mechanistic competence, and the robustness of the tier call across the Boltz-2 model ensemble. **(a)** Composition of the confidence-reactivity conflict classes. **(b)** Model confidence against the backside S_N_2 attack angle across the library. **(c)** Multi-model competence consensus by tier. **(d)** Model agreement across the five-model ensemble. **(e)** Catalytic competence against model confidence (Spearman ρ = +0.08) and against the mechanistic score (ρ = +0.61): confidence does not predict competence, while the mechanistic score shares most of its geometric terms with the competence score and so agrees with it largely by construction. **(f, g)** Robustness of the reactive geometry across the model ensemble, **(f)** across all models and **(g)** for the confidence-90 subset. Source data in SS_04.

### 3.4 A Mechanism-First Hierarchy Resolves the Conflict Between Confidence and Reactive Geometry

We replaced confidence-ranking with a configuration-weighted mechanistic score that measures the geometry a backside S_-_2 defluorination actually requires. Each complex is scored on the aspartate-oxygen to α-carbon distance, the S_-_2 attack angle, the internal catalytic-triad relay distances, the bidentate carboxylate clamp and the halide-stabilising cradle, with a Šidák correction that accounts for the multiple equivalent C-F bonds a polyfluorinated carbon presents ^49^. Every threshold and weight was fixed a priori and applied unchanged across all 58,050 complexes.

Applied library-wide, this filter separates the library into competence tiers (Figure 4). Only 8 complexes reach the elite Tier_1A, 47 reach Tier_1B and 731 Tier_2A (Figure 2d, e); a large intermediate band of 12,954 complexes populates Tier_2B, and the remainder distribute across Tier_3 (30,347), Tier_4 (2,557) and a Tier_5_Decoy set of 11,406 (Table 1). Median descriptors differed across tiers, though not in a monotonic way. The median S_-_2 attack angle is 173.2° in Tier_1A, 168.1 in Tier_2A and 114.4 in the decoys (Figure 4b, c), though it is not strictly monotonic across the intermediate tiers (156.6 in Tier_2B, 164.2 in Tier_3); the median aspartate-nucleophile distance tightens from 2.9 to 3.5 Å over the same range; and the median active-site RMSD to the crystal reference rises from 0.24 Å in Tier_1A to 3.2 Å in the decoys (Figure 4d; feature correlations, the gating cross-tabulation and the mechanistic-gate intersection are in Supplementary Figure 2). Median effective mechanistic scores were 1.00 in Tier_1A, 0.80 in Tier_2A, 0.39 in Tier_3 and 0.25 in Tier_5_Decoy. These separations follow the thresholds that define the tiers and are reported to characterise the resulting distribution, not as an independent test, while confidence and competence were weakly correlated (ρ = 0.08).

**Figure 4.**
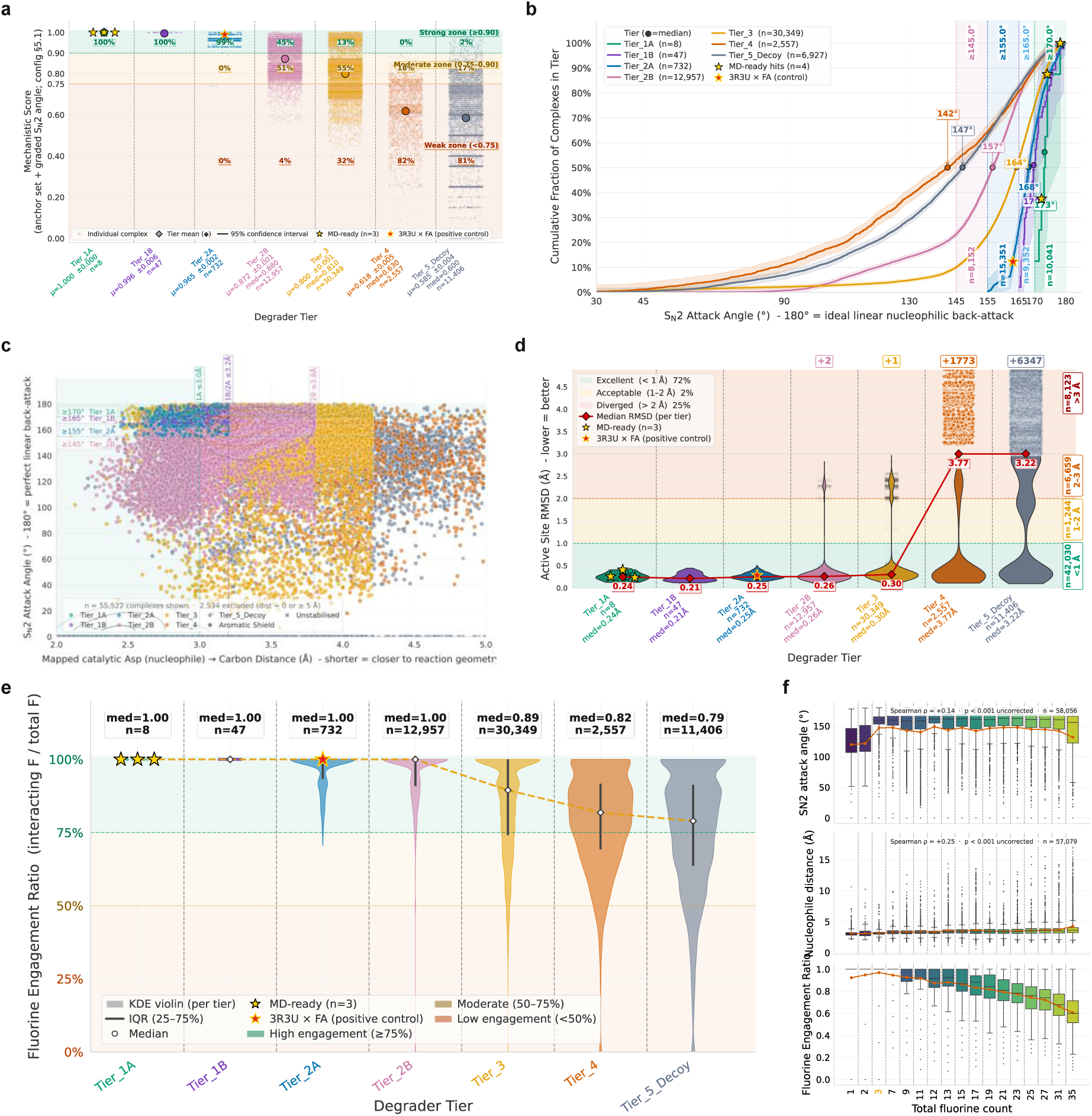
Catalytic mechanism and S_N_2 geometry. The mechanism-first score and the reactive geometry it measures. **(a)** Distribution of the configuration-weighted mechanistic score by tier. **(b)** Backside S_N_2 attack-angle distribution by tier, sharpening towards the ideal 180° in the elite tiers. **(c)** Reactive-centre geometry, the aspartate-nucleophile to α-carbon distance against the S_N_2 attack angle, coloured by tier. **(d)** Active-site RMSD to the crystal reference by tier. **(e)** Fraction of ligand fluorine atoms engaged in the reactive pocket, by tier. **(f)** Component breakdown of the mechanistic score by tier. Cross-tier statistics in SS_05; further diagnostics in Supplementary Figure 2.

**Table 1.** Mechanism-first competence hierarchy across the 58,050-complex FAcD-PFAS screen (2,150 variants against 27 substrates; the RPA1163/3R3U reference control is reported in Table 2; all six calibration controls are tabulated in SS_02). Each tier is summarised by its population and by the median of four independent mechanistic descriptors: the backside S_N_2 attack angle, the aspartate-nucleophile to α-carbon distance, the configuration-weighted mechanistic score (F4) and the active-site RMSD to the 3R3U crystal reference. The broadly monotonic sharpening of the composite mechanistic score from the decoy set to the elite tier (individual descriptors vary, particularly at Tier_2B), set against near-constant Boltz confidence (median 0.93–0.96 throughout), is the quantitative signature of the confidence-competence dissociation. Source data in SS_02.

| Tier | N complexes | % of library | Median $S_N2$ angle (deg) | Median Oδ-Ca (Å) | Median mechanistic score | Median active-site RMSD (Å) |
| --- | --- | --- | --- | --- | --- | --- |
| Tier_1A | 8 | 0.01 | 173.2 | 2.91 | 1.00 | 0.24 |
| Tier_1B | 47 | 0.08 | 169.7 | 3.05 | 1.00 | 0.21 |
| Tier_2A | 731 | 1.26 | 168.1 | 2.94 | 0.80 | 0.25 |
| Tier_2B | 12,954 | 22.32 | 156.6 | 3.16 | 0.69 | 0.26 |
| Tier_3 | 30,347 | 52.28 | 164.2 | 3.39 | 0.39 | 0.30 |
| Tier_4 | 2,557 | 4.40 | 142.2 | 4.04 | 0.24 | 3.77 |
| Tier_5_Decoys | 11,406 | 19.65 | 114.4 | 3.46 | 0.25 | 3.22 |

Decomposing the score into its individual mechanistic gates as an intersection (UpSet) analysis ^74^ showed that 17,838 complexes (30.7% of the library) satisfy every gate simultaneously, yet clearing all gates does not by itself earn elite status. The graded, penalty-adjusted score further separates the fully gated set, distributing these complexes across tiers, with only a handful reaching Tier_1A (Figure 4a), and the per-tier breakdown of its six components is shown in Figure 4f. The attack angle alone illustrates the same gate, though this follows from the conjunctive tier conditions rather than testing them. For fluoroacetate, difluoroacetate and trifluoroacetate, the library’s widest backside attack angle belongs in each case not to an elite complex but to one the pipeline classified as a lower likelihood of degradation: 177.9° and 179.7° in Tier_4 complexes for fluoroacetate and trifluoroacetate, and 179.2° in a Tier_5_Decoy complex for difluoroacetate, each closer to the ideal 180° than any elite complex with the same substrate attains. Among the ten widest angles per substrate, the tier assignments span three, four, and six of the seven tiers, respectively, covering every tier except Tier_2A.

All 55 complexes in the elite Tier_1A and Tier_1B are with the mono-, di- and trifluoroacetate ligands, showing a marked preference of FAcD enzymes toward ultra-short chain fluorinated substrates. Tier_1A is filled by five fluoroacetate, two difluoroacetate and one trifluoroacetate complexes, and Tier_1B by thirty fluoroacetate, fifteen difluoroacetate and two trifluoroacetate complexes, with best backside attack angles of 175.7, 177.1 and 179.3° respectively, each within 5° of the ideal 180° (across all elite complexes; the three named leads are in Table 2). These elite complexes are compact (median five ligand heavy atoms and one to two fluorines) and fully engaged, with every ligand fluorine contacted and both the halide-stabilising cradle and the bidentate carboxylate clamp present in all 55 cases; the long-chain perfluoroalkyl substances, whose reactivity is diluted over fifteen or more fluorines and a median twenty-six heavy atoms, fall to Tier_3 and below. The pipeline therefore recovers, without explicit instruction, the chemical logic of the fluoroacetate-dehalogenase family established by crystallography and kinetics: a backside S_-_2 defluorinase accommodates small, single-reactive-centre acids, and the screen concentrates fluoro-, difluoro-, and trifluoroacetate at the top ^8,11^. The 55 elite complexes derive from as many distinct sequences, a sequence-wide rather than single-lineage signal, and 157 variants place two or more substrates in the top three tiers, a one-enzyme-several-substrates breadth of practical relevance to a biocatalyst.

**Table 2.**
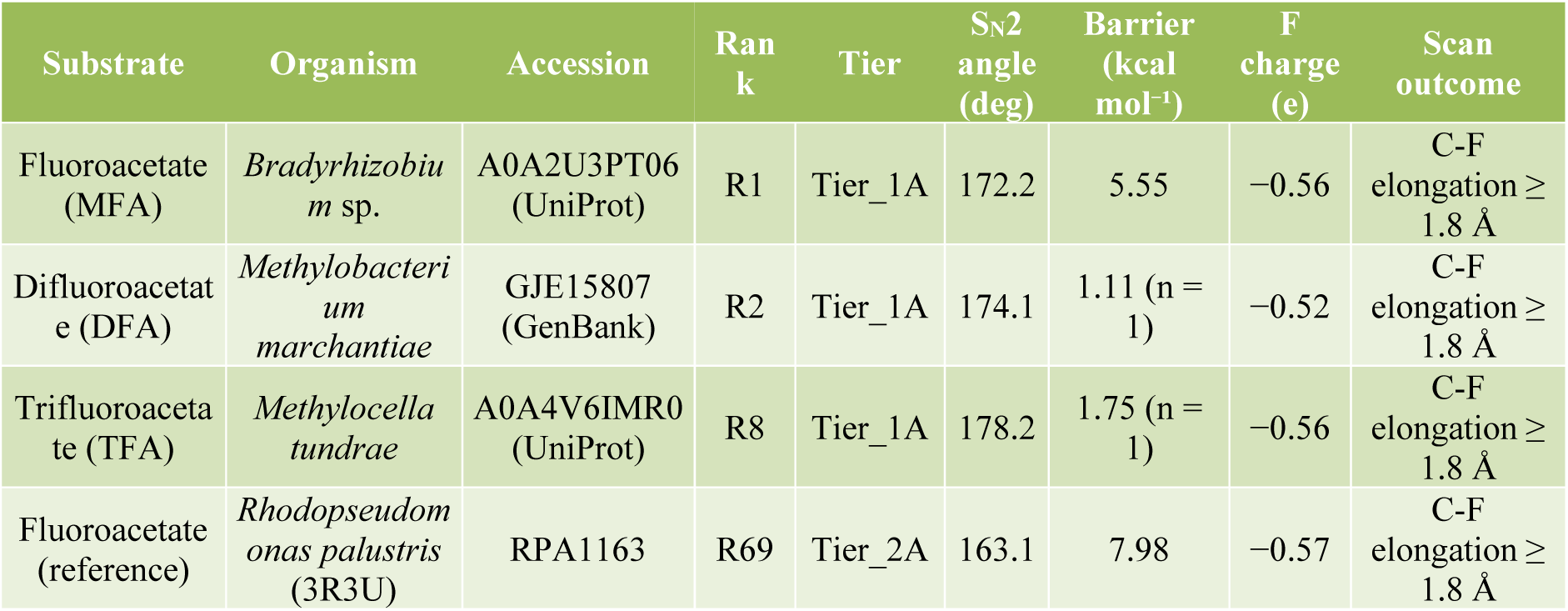
Top-ranked enzyme-PFAS pairs and the RPA1163/3R3U reference. The three variants that reach the elite Tier_1A against fluoro-, difluoro- and trifluoroacetate, with the crystallographic RPA1163/3R3U enzyme on its native substrate as the calibrating reference. The S_N_2 attack angle reflects the backside-attack geometry of the co-folded complex. The QM/MM barrier is a one-dimensional relaxed-scan maximum on the reactant side, is not transition-state-verified and is highest for the validated reference, so it does not rank the systems. The fluoride charge is the product-end departing-fluoride Mulliken charge, which corroborates but does not define the call. The turnover verdict keys only on the geometric criterion of F10 (product-end scissile C-F length ≥ 1.8 Å); it states that the active-site geometry permits cleavage, not that turnover is higher than the reference. Of the four substrates, only trifluoroacetate is a PFAS under the OECD definition. Per-system physics and QM/MM values are given in Table 3; source data in SS_02 and SS_14.

| Substrate | Organism | Accession | Rank | Tier | S<sub>N</sub>2 angle (deg) | Barrier (kcal mol<sup>-1</sup>) | F charge (e) | Scan outcome |
| --- | --- | --- | --- | --- | --- | --- | --- | --- |
| Fluoroacetate (MFA) | *Bradyrhizobium* sp. | A0A2U3PT06 (UniProt) | R1 | Tier_1A | 172.2 | 5.55 | -0.56 | C-F elongation ≥ 1.8 Å |
| Difluoroacetate (DFA) | *Methylobacterium marchantiae* | GJE15807 (GenBank) | R2 | Tier_1A | 174.1 | 1.11 (n = 1) | -0.52 | C-F elongation ≥ 1.8 Å |
| Trifluoroacetate (TFA) | *Methylocella tundrae* | A0A4V6IMR0 (UniProt) | R8 | Tier_1A | 178.2 | 1.75 (n = 1) | -0.56 | C-F elongation ≥ 1.8 Å |
| Fluoroacetate (reference) | *Rhodospseudomonas palustris* (3R3U) | RPA1163 | R69 | Tier_2A | 163.1 | 7.98 | -0.57 | C-F elongation ≥ 1.8 Å |

Across the 27-substrate panel, the fraction of the 2,150 variants reaching a degrader tier ranged from 78.0% for difluoroacetate and 77.8% for fluoroacetate down to zero for ten of the 27 substrates, comprising all six perfluoroalkyl sulfonates, both ether replacements (GenX and ADONA) and both fluorotelomer alcohols (Figure 5a). The Tier_1A enzyme-by-PFAS matrix locates the three leads within the elite set (Figure 5b). Competence is not confined to single pairings: 1,682 of the 2,150 variants reach a degrader tier against at least one substrate and 1,674 of these against two or more, with the twenty-five most promiscuous variants shown in Figure 5e, while within the top three tiers 157 of 572 variants engage two or more substrates. Pocket-residue reaction readiness tracks tier assignment across the panel (Figure 5f), and the radar profiles of the top hits and per-tier representatives, along with the ligand-to-tier circos map, summarise the same structure (Figure 5c, d, g).

**Figure 5.**
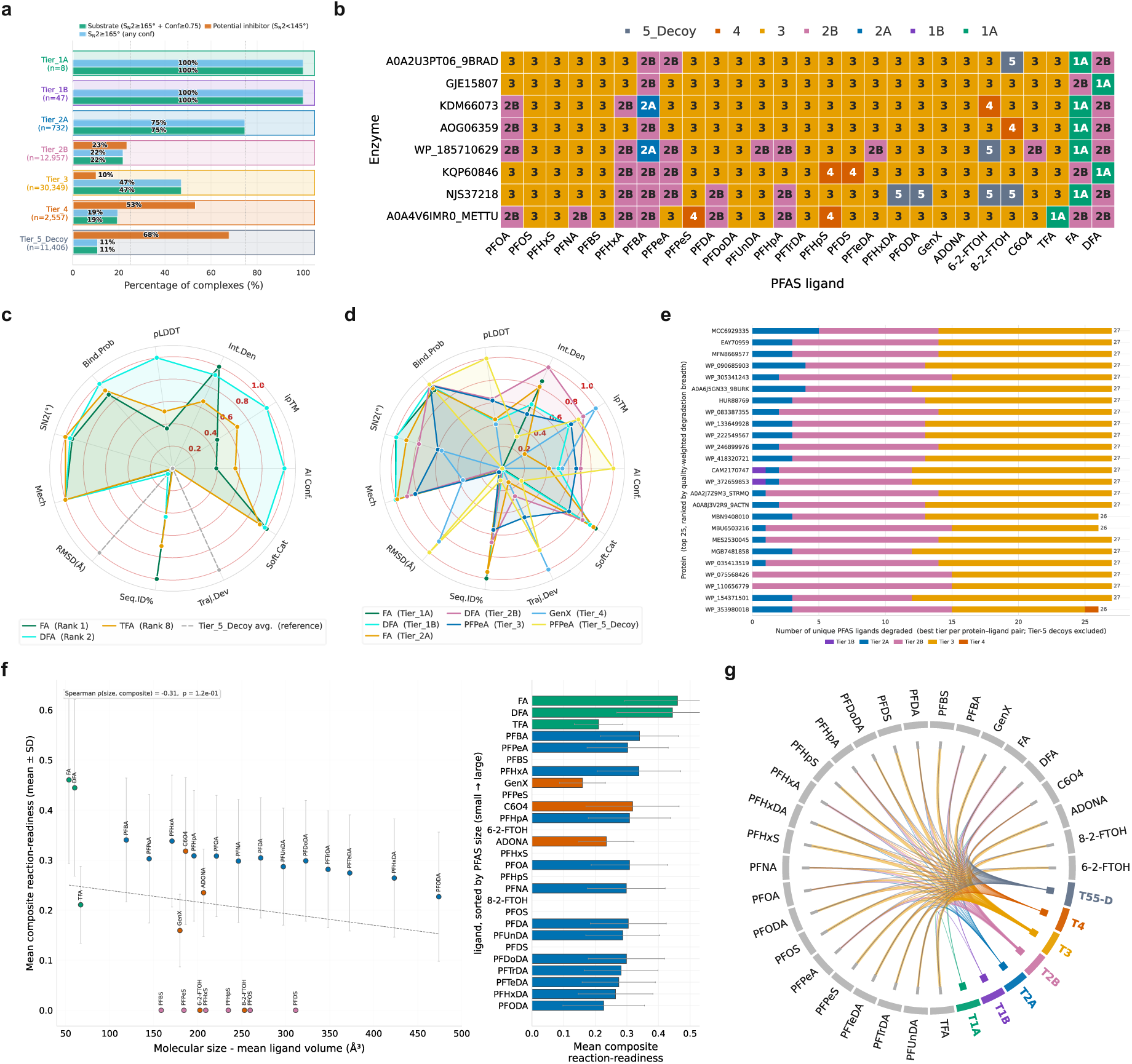
PFAS scope and lead candidates. Substrate scope and the leading enzyme-PFAS pairs. **(a)** Per-tier success rate across the PFAS panel. **(b)** Tier_1A enzyme-by-PFAS cross-ligand heatmap, locating the leads for fluoro-, difluoro- and trifluoroacetate (Table 2). **(c)** Radar profile of the top hits and **(d)** of the per-tier representatives. **(e)** Twenty-five most multi-target proteins. **(f)** Pocket-residue reaction-readiness map. **(g)** Circos map of ligand-to-tier assignment. The full ranked table is in SS_02; further scope diagnostics in Supplementary Figure 1.

The most consequential outcome is that the three top-ranked identified leads reach Tier_1A against the three benchmark substrates of increasing difficulty (Table 2): a *Bradyrhizobium* sp. variant (UniProt A0A2U3PT06) against fluoroacetate (attack angle 172.2°), a *Methylobacterium marchantiae* variant (GenBank GJE15807) against difluoroacetate (174.1°) and a *Methylocella tundrae* variant (UniProt A0A4V6IMR0) against trifluoroacetate (178.2°; recovered in one of five diffusion samples, see Limitations), all members of the Alphaproteobacteria. The trifluoroacetate lead had a near-ideal predicted backside attack angle (178.2°), identifying a natural variant for experimental testing and potential engineering ^23^ (Figure 5). Fluoroacetate dehalogenase is generally reported to be recalcitrant to trifluoroacetate ^17^, and the prediction made here is geometric and deliberately narrow: it states that the active-site geometry permits backside attack, not that turnover exceeds the reference, so it is a testable hypothesis rather than a measured rate.

Beyond the core S_-_2 geometry, the elite complexes engage a larger fraction of the ligand fluorine atoms, a median of 1.00 against 0.79 in the decoy set (Figure 4e), and present a distinct protein-ligand interaction fingerprint (Figure 6f–i) at a median interaction density of 4.0 against 1.5 interactions per ligand heavy atom (Supplementary Figure 3). The carboxylate head is anchored by a bidentate arginine clamp while the scissile fluorine is oriented into the aromatic and polar cradle, and the hydrogen-bond and halogen-contact geometry of these interactions is consistent with the criteria established for biological halogen bonding and fluoride recognition ^44^. The chemical-space map of the panel confirms that competent pairs are not confined to the smallest substrates but track the reactive-centre chemistry across the ligand set.

**Figure 6.**
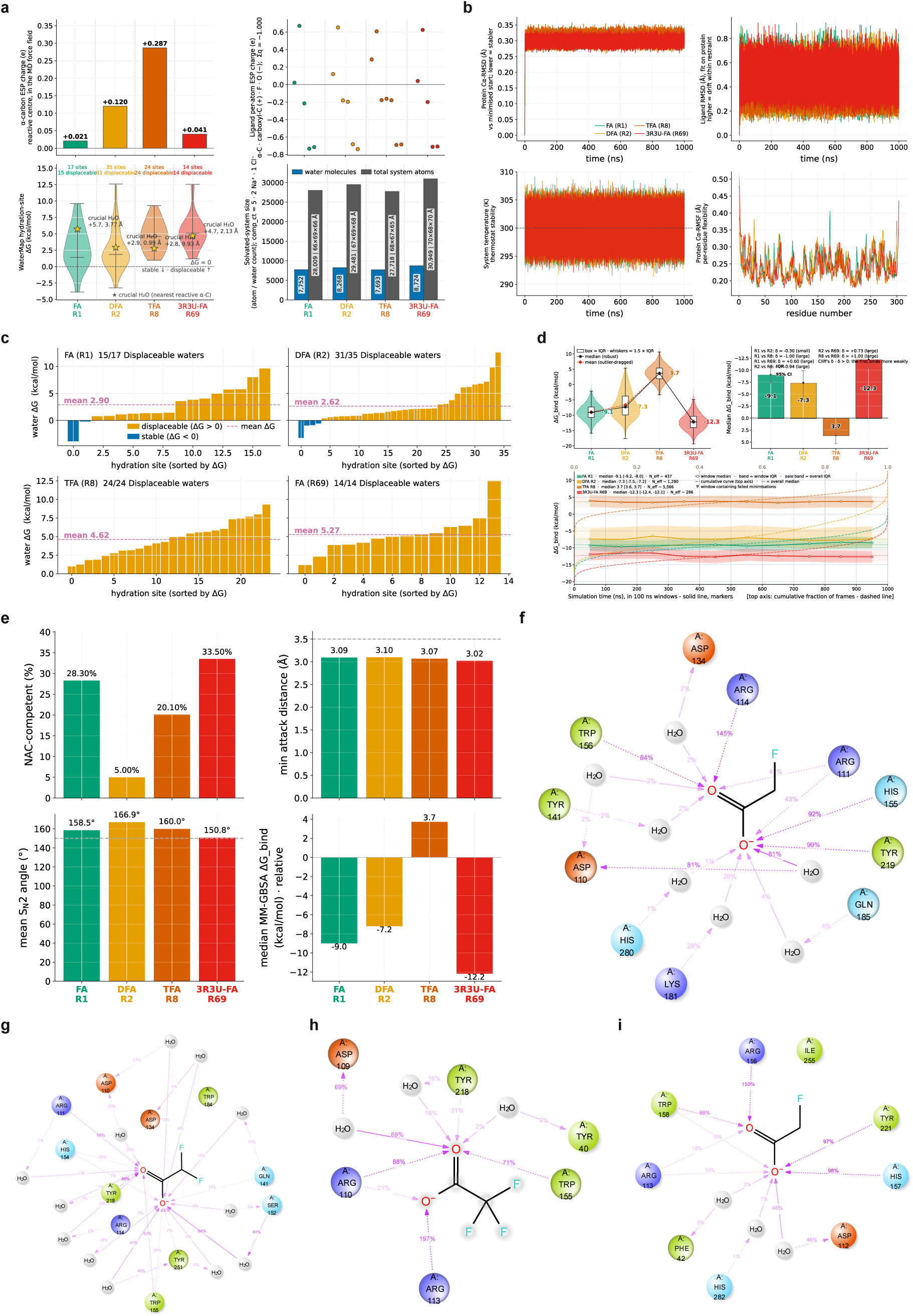
Physics, MD simulations and hydration. The physics-validation stages for the four selected systems, R1, R2, R8 and the R69 control. **(a)** Build and solvation quality control for the molecular-dynamics systems. **(b)** One-microsecond explicit-solvent trajectory quality control (protein and ligand RMSD, protein RMSF and temperature, all four ranks overlaid). **(c)** WaterMap hydration-site landscape across the four systems. **(d)** Per-rank MM-GBSA binding free energies computed with the VSGB 2.0 solvation model. **(e)** Cross-rank defluorination-geometry summary. **(f–i)** Active-site ligand-protein 2D interaction maps from the one-microsecond trajectories (contact persistence, percentage of frames, per residue): **(f)** R1, the fluoroacetate lead; **(g)** R2, the difluoroacetate lead; **(h)** R8, the trifluoroacetate lead; **(i)** R69, the RPA1163/3R3U control. Per-rank deep-dives are Supplementary Figures 6–9; the underlying values are in SS_08–SS_12.

### 3.5 The Screen Identifies Improved Attack Geometry and Reaches Challenging Substrates

A defining outcome of the screen is that its leading candidates are predicted to be more favourably poised, in backside-attack geometry, for defluorination than the reference enzyme itself. The crystallographic RPA1163/3R3U enzyme against its native substrate, fluoroacetate, is correctly recognised as catalytically competent and populates a high degrader tier (Tier_2A; S_-_2 attack angle 163.1°), validating that the mechanistic score identifies a bona fide defluorinase as engaged and calibrating the hierarchy against known chemistry. Ranked above it are natural variants promoted to the elite Tier_1A with markedly more ideal backside geometry (attack angles 172 to 178°). The pipeline therefore re-finds the known enzyme and also identifies sequence-diverse candidates with a more ideal backside attack geometry than the crystallographic reference enzyme for fluoroacetate, while simultaneously identifying variants already suited to difluoroacetate and for trifluoroacetate, the substrate the wild-type enzyme handles least well. The controls serve two purposes. They show that the score ranks the one experimentally validated enzyme highly, and they set the benchmark against which the new variants will be measured in the laboratory. The superiority established here is one of static reactive geometry alone; whether it converts into competitive turnover is not settled by that geometry, as the reference retains the highest near-attack dwell (below), and it awaits the quantum cleavage evidence and then experiment.

Three columns of Table 2 are QM/MM read-outs, described in full in section 3.8. Barrier (kcal mol⁻¹) is the maximum of a one-dimensional relaxed scan on the reactant side of the cleavage point, not a transition-state-verified activation energy. F charge (e) is the Mulliken charge on the departing fluoride; it supports the cleavage call but does not define it. Scan outcome is the call itself, made on geometry alone: the product-end scissile C-F length must reach 1.8 Å (Figure 7; Table 3).

**Figure 7.**
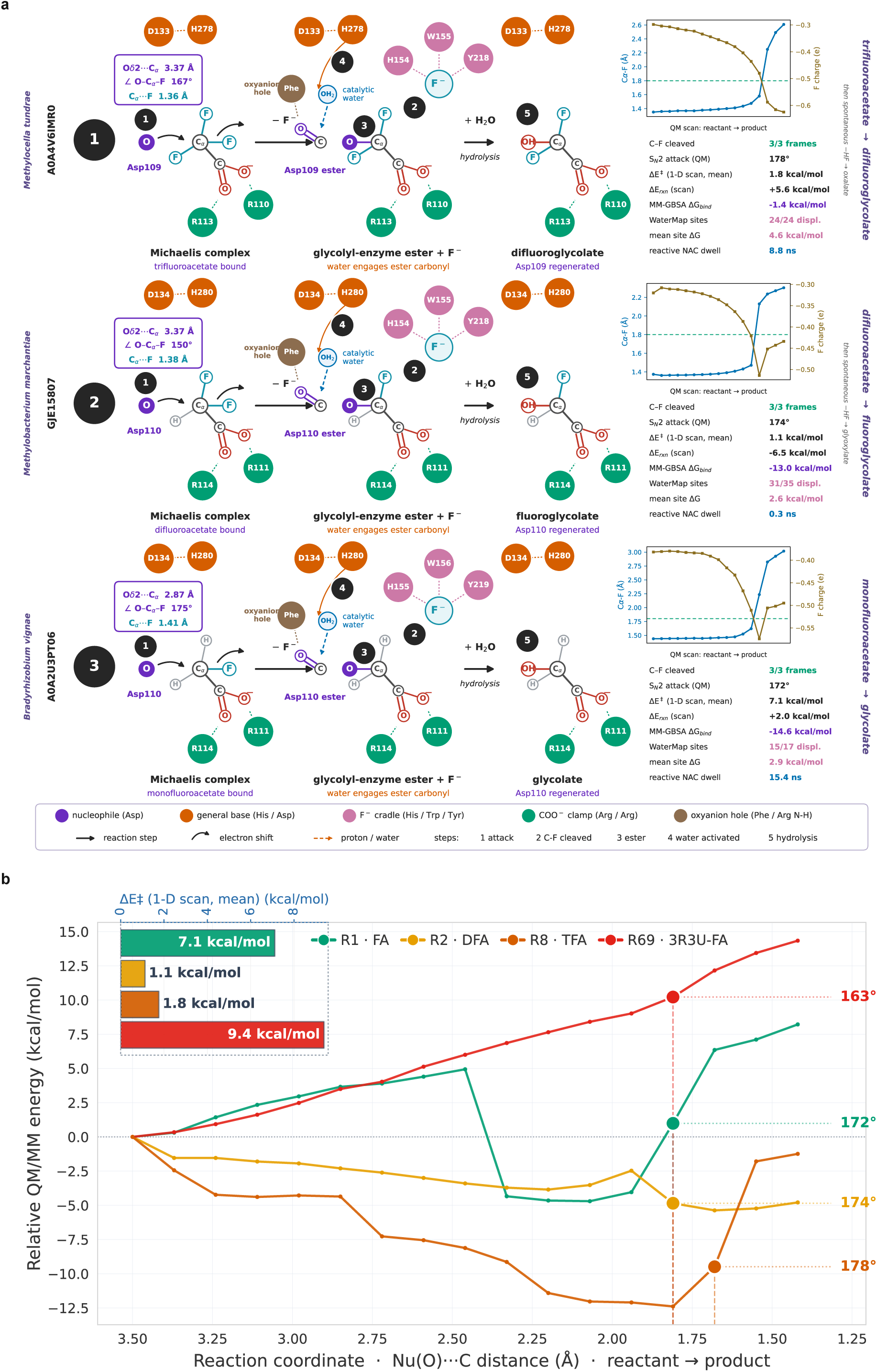
Enzymatic defluorination mechanism and QM/MM validation. **(a)** The hydrolytic double-displacement mechanism (Asp110 backside S_N_2 `→` glycolyl-enzyme ester `→` His280/water hydrolysis), drawn from the real QM geometry for the three independent lead FAcD variants, each on its own substrate (trifluoroacetate, Methylocella tundrae; difluoroacetate, Methylobacterium marchantiae; fluoroacetate, Bradyrhizobium sp.), with per-enzyme QM/MM reaction-coordinate plots (scissile C-F length and departing-fluoride charge) and a joined QM/MM-geometry-WaterMap-MD read-out; the enzyme replaces one C-F with C-OH, and for the di- and mono-fluoro α-hydroxy products a non-enzymatic HF elimination follows. These are three independent enzymes, one per substrate, not a single-substrate cascade. **(b)** All-jobs QM/MM potential-energy-surface overlay over the scissile C-F bond length, with the geometric cleavage verdict per system. Per-system relaxed-scan reaction profiles, including the RPA1163/3R3U reference, are in panel (h) of the per-system near-attack supplementary figures (Supplementary Figures 6–9).

**Table 3.**
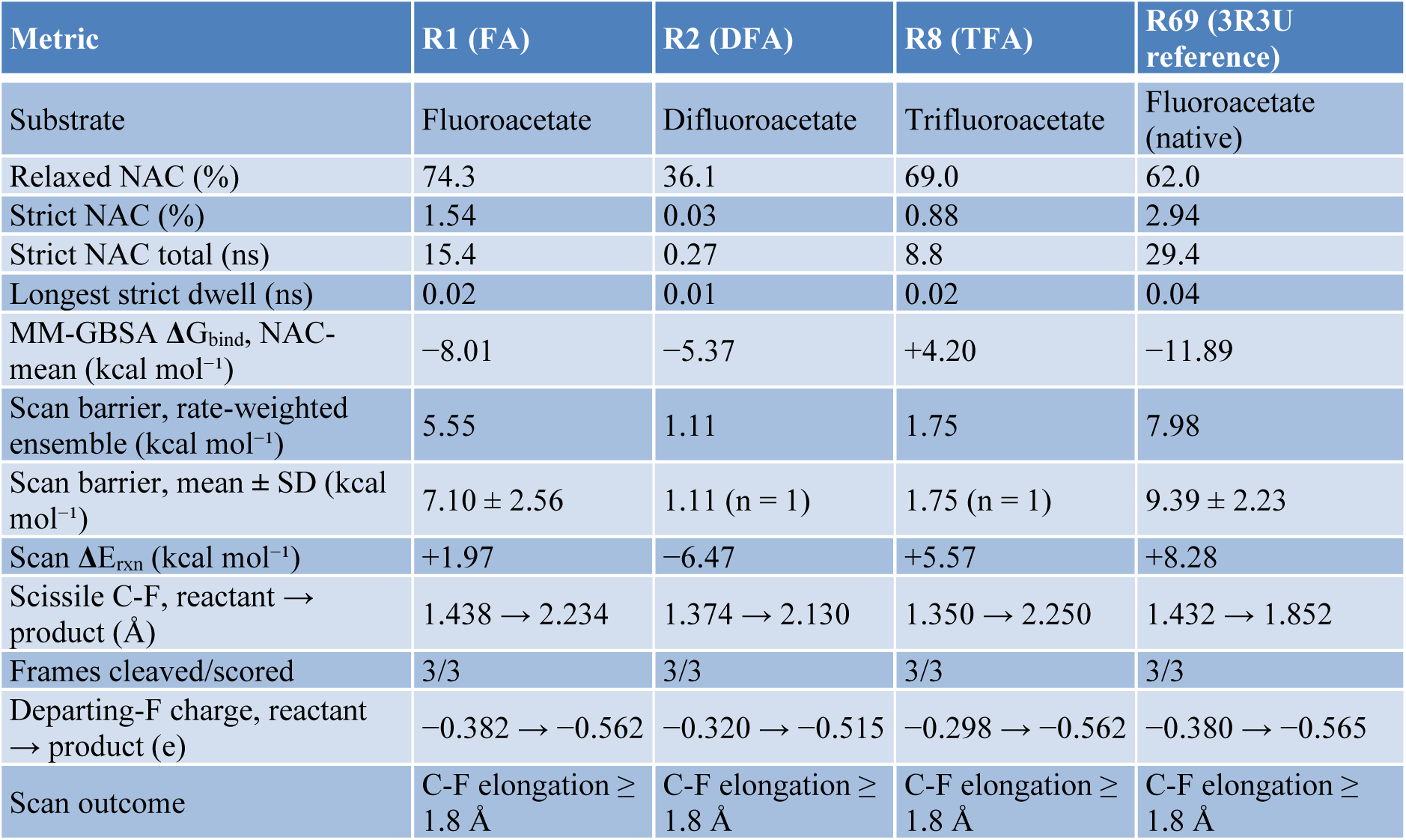
Physics and QM/MM characterisation of the four simulated systems. Near-attack statistics from the one-microsecond trajectories, endpoint MM-GBSA binding free energies and the QM/MM relaxed-scan read-outs for the three leads and the RPA1163/3R3U reference. The scan barrier is a one-dimensional relaxed-scan maximum on the reactant side, is not transition-state-verified, is not physically interpretable below about 1 kcal mol⁻¹, and is highest for the validated reference, so it does not rank the systems; the frame-to-frame standard deviation is comparable to the between-system spread. The verdict keys only on the geometric criterion of F10 (product-end scissile C-F length ≥ 1.8 Å). The ensemble barrier is the rate-weighted estimator of SF16, computed over the frames that resolved a transition state within the scan window; it is dominated by the lowest such barrier and is therefore not the arithmetic mean, which is reported separately. The trifluoroacetate lead (R8) binds unfavourably, as indicated by the endpoint estimate (Figure 6d). The NAC-mean is averaged over the near-attack frames of each trajectory, 141, 2, 97 and 299 frames for R1, R2, R8 and R69 respectively, so the R2 value rests on a small sample and is indicative only; it is not used to rank the systems, and the tier and cleavage calls for R2 do not depend on it. Of the four substrates, only trifluoroacetate is a PFAS under the OECD definition. Source data in SS_10–SS_14; the NAC-mean MM-GBSA values are the MMGBSA_dG_NAC_Mean_kcal column of the MD_Master_Ranking block in SS_13, which SS_11 does not carry.

| Metric | R1 (FA) | R2 (DFA) | R8 (TFA) | R69 (3R3U reference) |
| --- | --- | --- | --- | --- |
| Substrate | Fluoroacetate | Difluoroacetate | Trifluoroacetate | Fluoroacetate (native) |
| Relaxed NAC (%) | 74.3 | 36.1 | 69.0 | 62.0 |
| Strict NAC (%) | 1.54 | 0.03 | 0.88 | 2.94 |
| Strict NAC total (ns) | 15.4 | 0.27 | 8.8 | 29.4 |
| Longest strict dwell (ns) | 0.02 | 0.01 | 0.02 | 0.04 |
| MM-GBSA $\Delta G_{\text{bind}}$, NAC-mean (kcal mol<sup>-1</sup>) | -8.01 | -5.37 | +4.20 | -11.89 |
| Scan barrier, rate-weighted ensemble (kcal mol<sup>-1</sup>) | 5.55 | 1.11 | 1.75 | 7.98 |
| Scan barrier, mean ± SD (kcal mol<sup>-1</sup>) | 7.10 ± 2.56 | 1.11 (n = 1) | 1.75 (n = 1) | 9.39 ± 2.23 |
| Scan $\Delta E_{\text{rxn}}$ (kcal mol<sup>-1</sup>) | +1.97 | -6.47 | +5.57 | +8.28 |
| Scissile C-F, reactant → product (Å) | 1.438 → 2.234 | 1.374 → 2.130 | 1.350 → 2.250 | 1.432 → 1.852 |
| Frames cleaved/scored | 3/3 | 3/3 | 3/3 | 3/3 |
| Departing-F charge, reactant → product (e) | -0.382 → -0.562 | -0.320 → -0.515 | -0.298 → -0.562 | -0.380 → -0.565 |
| Scan outcome | C-F elongation ≥ 1.8 Å | C-F elongation ≥ 1.8 Å | C-F elongation ≥ 1.8 Å | C-F elongation ≥ 1.8 Å |

### 3.6 The Catalytic Triad and Halide Cradle are Temporally Stable Under Explicit-Solvent Dynamics

Static co-folded geometry indicates catalysis is possible but does not establish it, so we advanced the four highest-priority systems (ranks R1, R2 and R8, plus the R69 3R3U control) to one-microsecond explicit-solvent Desmond-based MD Simulations under the OPLS4 force field ^62,63^. Across the trajectories, the catalytic triad and the halide-cradle anchors were maintained, and the prepared reactive pose was retained throughout production. Because the ligand and backbone were positionally restrained, this retention is restraint-enforced and is not evidence that the co-folded geometry is intrinsically stable in solvent; what it establishes is that the catalytic machinery remains assembled around a held substrate, which is the precondition for the near-attack and cleavage analyses that follow (Figure 6b; Supplementary Figures 6–9, one per simulated system: R1, R2, R8 and the RPA1163 reference respectively; SS_12).

Every selected system sustained a measurable near-attack population over its one-microsecond trajectory, though the dwell is brief in absolute terms. The aspartate nucleophile and substrate backside geometry occupied the relaxed near-attack window in 74.3% of frames for the fluoroacetate lead, 69.0% for the trifluoroacetate lead and 36.1% for the difluoroacetate lead, compared with 62.0% for the RPA1163 reference on its native substrate, fluoroacetate. The tight strict near-attack criterion was met far less often (1.5, 0.9, 0.03 and 2.9% respectively), and the longest continuous strict residence never exceeded 0.04 ns in any system. As expected for a native enzyme acting on its physiological substrate, the reference accumulates the most strict near-attack time (29.4 ns over the trajectory); the leads, selected on static poise, not on dynamics, populate the strict window less but establish the dynamical prerequisite for defluorination that the co-fold alone cannot demonstrate. The difluoroacetate lead is the informative outlier: it enters the near-attack window least often yet, when it does, presents the tightest backside geometry (mean in-window attack angle 166.8°, against 158.4 and 160.2° for the fluoro- and trifluoroacetate leads), consistent with a tightly packed active site that samples fewer but geometrically cleaner attack conformations. Because the near-attack dwell is low across every system, including the validated reference, it constrains but does not by itself decide turnover, which is adjudicated by the geometric C-F cleavage below (Figure 6b, e; Figure 7b; Table 3; Supplementary Figures 6–9, panel (h); SS_13).

### 3.7 Hydration of the Reactive Pocket Differs Across the Selected Systems

WaterMap hydration analysis located the structural and displaceable hydration sites of the reactive pocket in the ligand-bound (holo) state ^67^. Displacement of energetically unfavourable waters accompanies binding for the fluoroacetate and difluoroacetate leads, whose reactive pockets carry thermodynamically stable hydration sites; the trifluoroacetate lead enzyme and the RPA1163 reference carry none, so no such displacement term applies to them (Figure 6c; stable-water counts: R1 two, R2 four, R8 zero, R69 zero). The NAC-conditioned endpoint MM-GBSA binding free energies are favourable for the fluoroacetate and difluoroacetate leads and for the reference (−8.0, −5.4 and −11.9 kcal mol⁻¹, respectively, computed with the VSGB 2.0 solvation model) ^68,69^. The trifluoroacetate lead is the exception: its endpoint MM-GBSA value is unfavourable (+4.2 kcal mol⁻¹, NAC-conditioned), so its predicted binding is weak, and its case rests on reactive geometry and geometric cleavage, not on affinity. These endpoint estimates are reported for context, not as a ranking criterion (Figure 6d; hydration-site thermodynamics in SS_10, MM-GBSA decomposition in SS_11).

### 3.8 QM/MM Relaxed Scans of the C-F Cleavage Coordinate

The scanned geometries reproduce the first displacement of the hydrolytic double-displacement mechanism in all three leads (Figure 7a). In each, the aspartate nucleophile attacks Cα from the backside and displaces fluoride into the halide cradle, giving the glycolyl-enzyme ester, which the histidine base is expected to hydrolyse, regenerating the aspartate; that hydrolytic step is taken from the established mechanism rather than computed here. Because the nucleophile is an aspartate carboxylate oxygen, the enzyme substitutes C-OH for C-F rather than C-H, so the expected primary products are the corresponding α-hydroxy acids: difluoroglycolate from trifluoroacetate, fluoroglycolate from difluoroacetate and glycolate from fluoroacetate. The two polyfluorinated products are gem-halohydrins and are expected to eliminate HF non-enzymatically, releasing a second fluoride, which is the route by which difluoroacetate is reported to yield glyoxylate 13. The scans resolve the first displacement through to ester formation; the hydrolysis and the HF elimination are not computed. The three systems are three independent enzymes acting on three different substrates, not one enzyme stepping a single substrate down a series. Catalytic roles were assigned in each variant by alignment to the RPA1163/3R3U reference, so the residue numbers in Figure 7a differ between rows and are each variant’s own.

The relaxed scans resolve C-F cleavage geometrically. In each lead the scissile C-F bond lengthens from about 1.4 Å at the reactant to 2.23 Å (fluoroacetate), 2.13 Å (difluoroacetate) and 2.25 Å (trifluoroacetate) at the product, and to 1.85 Å in the RPA1163 reference, past the 1.8 Å cleavage threshold in all three sampled reactive frames of every system; for the trifluoroacetate lead enzyme the product-end distance reached 2.11 to 2.29 Å across its three frames, every one of them above the threshold, and the Mulliken charge on the departing fluoride falls to between −0.52 and −0.57 e, a partial charge consistent with fluoride departure but short of a fully released ion: a free fluoride ion would carry close to −1 e, so a value near −0.55 e means the bond is geometrically broken while the departing fluorine still shares electron density with the carbon (the diffuse-function-free basis under-reports the free-ion charge, so the geometric criterion, robust to the basis, defines cleavage). The one-dimensional relaxed scan drives only the forming nucleophile-carbon coordinate, so the energy maximum on its reactant side is a scan feature rather than a transition-state-verified activation energy; and the apparent barrier is not monotonic in screen rank and is highest for the reference enzyme (7.98 kcal mol⁻¹, against 5.55 for the fluoroacetate lead and 1.11 from the single converged frame of the difluoroacetate lead; 1.75 from the single converged frame of the trifluoroacetate lead), which is why the turnover verdict is anchored to the geometric cleavage criterion, not to the scan barrier. Tables 2 and 3 report the barriers for context (Figure 7b; Supplementary Figures 6–9, panel (h); SS_14), read against the established backside-S_-_2 route of FAcD, defined by crystallography and reaction-coordinate analysis ^11^. Three near-attack frames were scanned per system, chosen as the most pre-organised frames of the trajectory, so the barrier carries a selection bias; the frame-to-frame standard deviation of the barrier, where enough frames converged to define one, is comparable to the difference between systems (7.10 ± 2.56 kcal mol⁻¹for the fluoroacetate lead and 9.39 ± 2.23 for the reference, each over three converged frames; the difluoro- and trifluoroacetate leads converged in one frame apiece and so support no spread). The scan reaction energy is downhill only for the difluoroacetate lead (−6.47 kcal mol⁻¹) and uphill for the fluoroacetate and trifluoroacetate leads and the reference (+1.97, +5.57 and +8.28 kcal mol⁻¹), and is likewise a scan feature. For the trifluoroacetate lead, the single converged frame gives a barrier of 1.75 kcal mol⁻¹, close to the floor below which a relaxed-scan maximum is not physically interpretable, so its cleavage call rests on the geometric criterion alone. Every system, including the positive control, cleaves in all three frames (3/3). The size of the quantum region follows the measured hydration, with first-shell waters selected within 3.5 Å of the reaction centre and capped at three: three entered for the trifluoroacetate lead and for the reference, two to three for the fluoroacetate lead, and two for the difluoroacetate lead. The first-shell solvation of the departing fluoride is therefore treated quantum-mechanically in every system (SS_10, SS_14).

On the geometric criterion, all three lead enzymes register as defluorinating (Figure 7b; Supplementary Figures 6–8, panel (h)), with C-F bond breaking in every sampled reactive frame, as does the RPA1163 reference on its native substrate (Supplementary Figure 9, panel (h)), confirming that the criterion recovers the one experimentally validated defluorinase in the set. Three caveats bound what this step establishes, and they apply to all four systems. First, the relaxed scan drives only the nucleophile-carbon coordinate, so the energy maximum on its reactant side is a scan feature rather than a transition-state-verified activation energy, and the elongation past the cleavage threshold is in part a mechanical consequence of that drive. Second, the apparent barrier is not monotonic in screen rank and is highest for the reference enzyme, and the near-attack dwell is low in every system, so conditioning the verdict on either would reject the validated reference itself; both are reported as context rather than imposed as independent gates (Figure 8a-f; Table 3; SS_11, SS_12, SS_14). Third, because every system cleaves in all three frames, the set contains a positive control but no demonstrated negative. Discrimination in this study occurs upstream, at library scale, though that separation follows from the thresholds that define the tiers and characterises the resulting distribution rather than testing it, since no low-scoring complex was carried to QM/MM: the mechanistic score separates all 58,050 complexes on backside-S_-_2 geometry, and the Tier_5_Decoy complexes, whose median S_-_2 attack angle is 114.4° versus 173.2° in Tier_1A, are the geometric negatives. The QM/MM scan therefore confirms the single pre-selected elite winner per substrate, not the discriminator.

**Figure 8.**
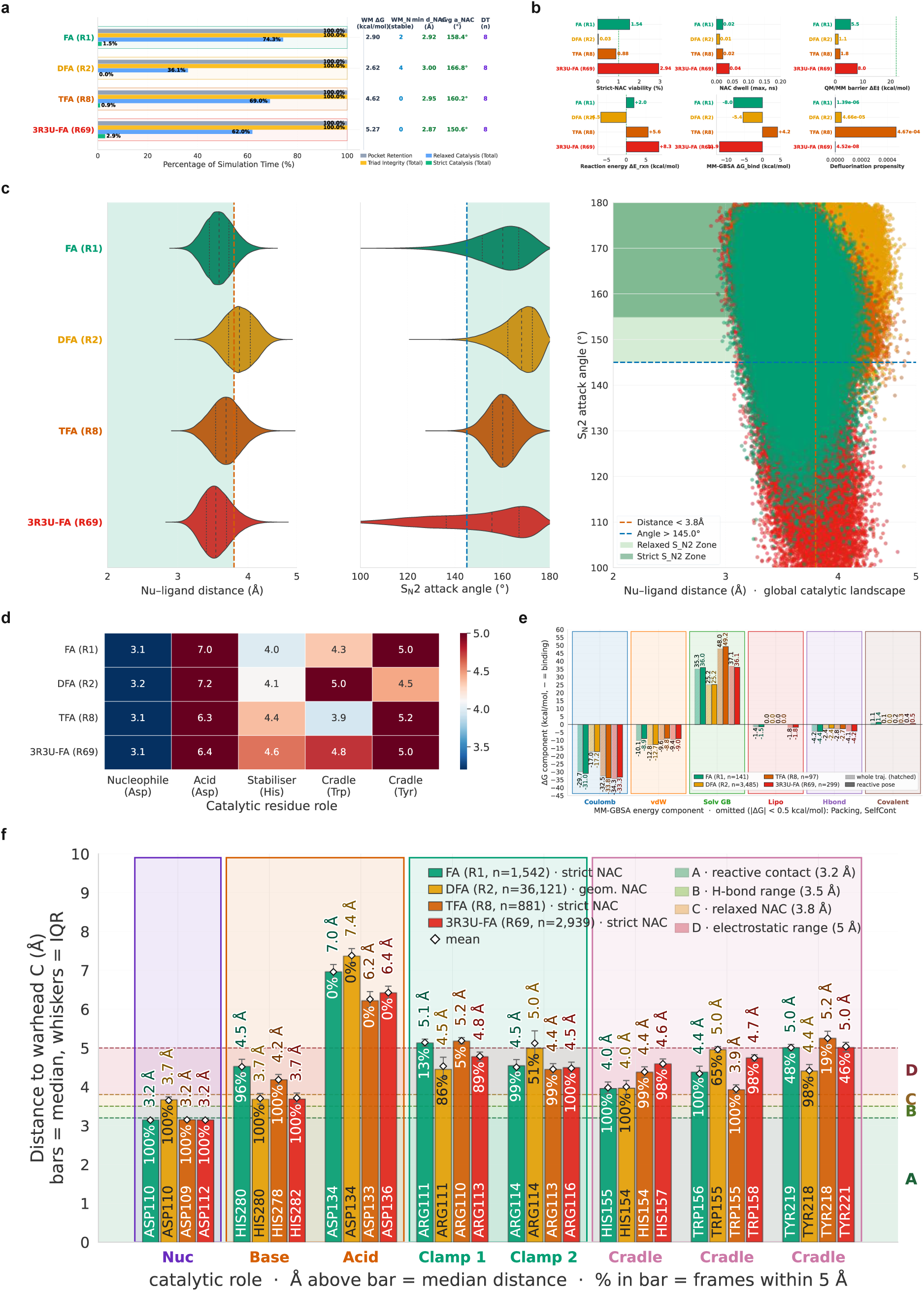
Cross-rank defluorination outcome. Cross-rank summary of the QM/MM defluorination stage across the four scanned systems (R1, R2, R8 and the R69 control). **(a)** Viability summary. **(b)** Defluorination landscape. **(c)** Comparative analysis of the QM/MM, geometry and hydration read-outs. **(d)** Comparative active-site residue engagement across the systems. **(e)** MM-GBSA energy-term decomposition across ranks. **(f)** Catalytic-machinery engagement across ranks. Per-system scans are panel (h) of Supplementary Figures 6–9; source data in SS_14.

## 4. Discussion

The central methodological result of this study is a clean dissociation between artificial-intelligence confidence and catalytic competence. Boltz-2 assigned near-identical, uniformly high confidence (median 0.93 to 0.96) to complexes spanning the entire mechanistic spectrum, from elite, near-ideally poised poses to catalytically inert decoys, and that confidence carries almost no information about whether an enzymatic complex is catalytically poised. A screen that ranks by confidence, as co-folding pipelines commonly do, would treat the library as equivalent and yield nothing to select as a preferred complex for lab validation. This observation is not an idiosyncrasy of one model or one target: it is the direct, enzyme-specific manifestation of the emerging finding that deep-learning co-folding models do not reliably learn the physics of protein-ligand interaction and are biased toward the distribution of their training data ^29^. Imposing an explicit, configuration-defined mechanistic gate, keyed on the reactive geometry a backside S_-_2 defluorination actually requires, recovers a steep competence hierarchy in which only eight of 58,050 complexes reach the top Tier_1A. This is achieved by mechanistic gating, not the generative model. The caution for any catalytic use of co-folding follows directly: model confidence indicates whether a pose is well formed, not whether it can react, and this physics-anchored filter converts a confident structural prediction into a testable catalytic hypothesis.

The discovery-level finding is that standing environmental sequence diversity already contains FAcD active-site geometries suited to poly- and perfluorinated substrates. Three natural enzyme variants reached the elite tier against fluoro-, difluoro- and trifluoroacetate, the prototypical Teflon-shielded PFAS target, with near-ideal backside attack angles. This complements the powerful protein-engineering campaigns now being reported for FAcD: the gate-based synergetic chain redesign of the RPA1163 scaffold delivered variants with very high turnover on a non-natural α-fluoro substrate, but proceeded from a single, well-characterised enzyme and a defined mutational logic ^23^. Screening natural FAcD variants by reactive geometry provides a shortlist for experimental testing and may inform future enzyme engineering.

Mechanistic knowledge, structural data and prior experimental findings support the plausibility of these predictions; enzyme activity in the identified variants remains to be tested. First, the mechanistic filter is not a free parameter fitted to our data: it encodes the catalytic geometry that crystallography, mutagenesis, and reaction-coordinate kinetics have established for FAcD, including the aspartate backside S_-_2 attack, the aspartate-histidine relay, and the halide-stabilising cradle. A complex scoring highly reproduces an experimentally validated active site, and Tier_1A recovers a median nucleophile distance of 2.9 Å and a median attack angle of 173.2°, matching the crystallographic reactive geometry ^8,11^. Second, the positive control ties the *in silico* scale to a real enzyme: the experimentally characterised RPA1163/3R3U fluoroacetate dehalogenase is correctly recognised as engaged and competent on its native substrate, so the hierarchy is calibrated against known chemistry rather than asserted ^17^. Third, the substrates our emergent screen promotes are precisely those for which enzymatic defluorination has been demonstrated in the laboratory: FA is the canonical FAcD substrate, and hydrolytic defluorination of difluoroacetate has been shown experimentally. Hence, the difluoroacetate-cleaving enzyme extends a demonstrated reaction rather than postulating a new one ^13^. The contribution of this work is a mechanism-anchored discovery pipeline rather than a single new enzyme or a new abiotic process: it takes a sequence database and a set of AI co-folds into an experimentally testable, geometry-ranked shortlist, and that recovers the known chemistry of the family as an internal control while extending it to the harder polyfluorinated acids.

Compared with current defluorination approaches, enzymatic routes offer the prospect of mild, selective, low-cost methods. DeFluorX provides a scalable framework to screen millions of enzymes against fluorinated substrates and prioritise candidates through reproducible, physics-based validation. Abiotic technologies, particularly recent photocatalytic systems that achieve near-complete or low-temperature PFAS mineralisation, are impressive but energy-intensive and non-selective, and are best suited to concentrated waste streams ^4,5^. Biological defluorination, by contrast, offers a mild, selective and in-principle scalable complement for the dispersed, low-concentration point sources that dominate real contamination, and the persistent bottleneck has been the scarcity of enzymes competent against the harder substrates^7^. By recasting that bottleneck as a searchable, mechanism-defined problem, this work provides a systematic front end for populating the enzymatic side of the remediation toolbox.

For the lead enzymes, the screen establishes better static reactive geometry. Its scope is limited as follows. The three Alphaproteobacterial variants present backside attack angles of 172 to 178° against the reference enzyme’s 163°, and the QM/MM scans show that their active-site geometry permits C-F cleavage, as it does for the validated reference. The screen does not establish higher turnover: the reference retains the highest near-attack dwell over the trajectory, the one-dimensional relaxed-scan barrier does not order the candidates and is highest for the reference, and the trifluoroacetate lead binds unfavourably by the endpoint estimate. The defensible claim is that natural diversity supplies variants geometrically better pre-organised than the crystallographic reference for the harder substrates, and that whether this pre-organisation converts into competitive catalysis is an experimental question. The screen narrows the sequence library to candidates for testing; assays are needed to determine whether their predicted geometries yield measurable turnover.

More generally, the architecture demonstrated here, layering an explicit mechanistic gate and a sequence of increasingly rigorous physics (MD Simulations, near-attack sampling, endpoint free energies, hydration thermodynamics and quantum mechanics) on top of AI co-folding, is not specific to FAcD. Any enzyme whose chemistry defines a measurable reactive geometry could be screened in the same way, and the approach directly addresses the physical realism gap that limits the use of co-folding predictions for catalytic questions ^29^. The near-attack-conformation formalism is central to this transferability because it connects a dynamically sampled geometry to the probability of barrier crossing and, therefore, to turnover ^19^.

In every lead complex, the scissile C-F bond broke in all sampled reactive frames, elongating past the 1.8 Å cleavage threshold with the departing fluoride reaching about −0.55 e; the same criterion recovers cleavage in the RPA1163 reference, confirming that it identifies the one experimentally validated defluorinase in the set. The one-dimensional relaxed-scan barriers are reported as context, not as transition-state-verified activation energies: because the scan drives only the forming nucleophile-carbon coordinate, the reference enzyme returns the highest apparent barrier, so the energetics cannot, by themselves, rank the leads, and the verdict is anchored to the geometric cleavage that all systems share. The discrimination in this work is made upstream, at library scale: the mechanistic score separates all 58,050 complexes on backside-S_-_2 geometry, with the Tier_5_Decoy complexes (median S_-_2 attack angle 114.4° against 173.2° in Tier_1A) as the geometric negatives, whereas the QM/MM scan confirms cleavage on the single pre-selected elite winner per substrate rather than acting as the discriminator. A one-dimensional driven scan cannot by itself separate mechanical from catalytic elongation, so the cleavage call is corroborated by that upstream geometric discrimination and by the recovery of the RPA1163 reference, an experimentally validated defluorinase, as a positive control. The established backside-S_N_2 mechanism of FAcD provides the reference reaction coordinate against which the scan maxima are read ^11,17,75^.

Several limitations bound the present claims and are noted here. The reproducibility of the elite assignment varies with substrate. The fluoroacetate and difluoroacetate leads were recovered by all five diffusion samples, whereas the trifluoroacetate lead, and the two other trifluoroacetate complexes in the elite tiers, were recovered by one sample of five. Trifluoroacetate assignments at the top of the ranking are therefore less reproducible than those for the mono-and difluorinated substrates, and the trifluoroacetate lead in particular should be treated as a single-sample observation pending experimental testing. The multiplicity correction also disadvantages polyfluorinated substrates relative to taking the best-aligned bond: it costs a monofluorinated carbon nothing, a difluorinated carbon a mean of 9.4° and a trifluoromethyl carbon a mean of 15.5°, although the best-aligned trifluoromethyl carbon in the library loses only 0.2°, so the penalty falls on angles that arise from having had several chances rather than on genuine alignment. The three leads clear the cleavage criterion by 0.03 to 0.49 Å, while the wild-type reference clears it by as little as 0.007 Å. The co-folded poses are static and inherit the training-data biases of the underlying model; we mitigate this with explicit-solvent MD Simulations, but the dynamics themselves rest on a fixed-charge force field, and the MM-GBSA ranking is an endpoint approximation rather than a rigorous free-energy calculation 68. The endpoint estimator was chosen for cost at this cohort size, 10,005 frame-level evaluations, and its values are used only as context alongside the geometric and QM/MM read-outs, never as a ranking criterion. The hydration analysis uses a rigid three-site water model whose self-diffusion is faster than experiment, so the reported water-residence signatures are conservative lower bounds. The quantum treatment uses the B3LYP functional with hydrogen-capped covalent cuts and an enforced neutral-histidine, deprotonated-aspartate active site; these are defensible, mechanism-motivated choices, but the protonation enforcement and the level of theory are assumptions, not measurements. Because the explicit quantum waters are chosen from the measured first shell within 3.5 Å of the reaction centre, the size of the quantum region is itself substrate-dependent, so the first-shell solvation of the departing fluoride is treated quantum-mechanically wherever such waters are present; the count is capped at three waters for tractability, and the remaining solvent is carried by the classical region. Above all, every result here is computational: the identified variants are predictions, and heterologous expression and fluoride-release assays must confirm their defluorination activity before any bioremediation claim can be made.

The immediate priority is experimental validation. The lead variants can be expressed and assayed for fluoride release; active candidates could then serve as starting points for engineering ^23^. The pipeline could also be adapted to related dehalogenase families and other enzymes with defined catalytic geometries.

## 5. Conclusion

**DeFluorX** integrates sequence, structural and mechanistic criteria to prioritise candidate fluorine-degrading enzymes from large sequence-substrate spaces. We co-folded 2,150 curated bacterial FAcD sequences against 27 environmentally prioritised fluorinated substrates using Boltz-2 and then ranked them mechanistically. Boltz-2 confidence scores varied little across the seven competence tiers (median, 0.929–0.955) and were only weakly associated with competence (ρ=0.08), providing limited differentiation among candidates. By contrast, the structural criteria separated competent complexes from decoys: the median attack angle decreased from 173.2° in Tier_1A to 114.4° in the decoy tier (ε² = 0.26). 30.7% of the library passed all criteria, indicating that these filters were selective without being overly restrictive. Together, these results show that model confidence alone does not reliably distinguish catalytically competent complexes, whereas mechanistic geometry provides a more informative basis for prioritising defluorinating candidates.

Eight of the 58,050 complexes reached Tier_1A, and 55 reached the two elite tiers, representing 55 distinct sequences. Variants from *Bradyrhizobium* sp., *Methylobacterium marchantiae* and *Methylocella tundrae* reached Tier_1A against fluoroacetate, difluoroacetate and trifluoroacetate, with attack angles of 172.2°, 174.1° and 178.2°, compared with 163.1° for the crystallographic RPA1163/3R3U reference. What distinguishes these candidates is reactive geometry, not demonstrated turnover. The reference retains the highest near-attack dwell and the highest relaxed-scan barrier, with the latter serving as a comparative descriptor across these four systems rather than a verified activation energy. In the quantum treatment, the scissile C-F bond elongated beyond 1.8 Å in every sampled frame of every system, and the departing fluoride reached about −0.55 e, supporting a reactive configuration but not establishing the rate or extent of cleavage. The top-ranked complexes were concentrated among the shorter fluorinated substrates, whereas longer-chain and ether-containing substrates were not represented in the elite tiers. This observation is consistent with the characterised FAcD enzymes but warrants more detailed investigation of substrate scope. Of the four systems carried to quantum treatment, only trifluoroacetate is a PFAS under the OECD definition. These results support mechanistic geometry as a useful basis for prioritising defluorinating candidates and identify the three leads as concrete, testable predictions for heterologous expression and fluoride-release assays against fluoroacetate, difluoroacetate and trifluoroacetate.

## Supporting information

https://doi.org/10.5281/zenodo.23318610

## Declarations

### Ethical responsibilities

not applicable; the study is computational.

### Consent for publication

all authors consent to submission.

## Acknowledgement

The authors thank the members of the Microbial Ecology and Biotechnology Section (Now Environmental Sciences).

## Data Availability

The data underlying all figures and tables are provided in the supplementary workbook Supplementary_Sheets_DeFluorX.xlsx, which is permanently archived on Zenodo (https://doi.org/10.5281/zenodo.23318610). The workbook also includes the source protein sequences and ligand structures used in this study. The Index sheet provides a mapping of each SS_nn identifier to its corresponding worksheet and contents.

## Code Availability

The DeFluorX pipeline code is available at https://github.com/KU-MGB/DeFluorX.

## Author Contributions

**SA** designed and implemented the pipeline, performed the analysis, wrote all the code, and wrote and revised the manuscript. **AF** contributed to the study design and provided experimental insight that informed pipeline development. **BT** contributed to methodology design and optimisation. **JBR** contributed to methodology design and optimisation. **MO** contributed to the study design and provided experimental insight that informed pipeline development. **TKN** secured funding, supervised the study, contributed to the study design, and critically revised the manuscript. All authors reviewed and approved the final manuscript.

