## Supplementary material for "Mechanism-guided Discovery of Fluoroacetate Dehalogenase (FAcD) Variants for Enzymatic PFAS Defluorination": https://doi.org/10.5281/zenodo.23318610

#### Scoring and Statistical Formulae:

This section gives the full mathematical definition of every scored quantity used to rank the 58,050 complexes and to validate the leads. Supplementary Table S1 lists all symbols, weights and thresholds, Supplementary Data SS_01 documents the sequence input and output provenance, and SS_02–SS_14 provide the per-complex values. Equations SF1–SF8, SF11, SF14 and SF15 are the expanded forms of main-text formulae F3–F13; SF9, SF10, SF12, SF13, SF16, SF17 and SF18 are defined only here.

**Mechanistic Score:** Each complex is scored on six reactive-geometry components combined using fixed weights that sum to one (SF1). Five are binary anchor checks, each contributing its full weight when its threshold is met: the nucleophile near-attack reach $g_{nuc}$ (Oδ–Cα ≤ 3.2 Å), the nucleophile-base relay $g_{NB}$ and base-acid relay $g_{BA}$ (≤ 5.0 and ≤ 5.5 Å), the bidentate carboxylate clamp $g_{clamp}$ and the halide-stabilising cradle $g_{stab}$. We established the residue identities and catalytic roles of these five through crystallography and mechanistic analysis [1,2], and the reach threshold follows the near-attack-conformation criteria of Lightstone and Bruice [3]. Only the Šidák multiplicity-corrected backside attack angle $g_{angle}$ is graded continuously [4]: it is the Šidák survival evaluated at the effective angle of SF2, $g_{angle} = [1 + cos(180° - \theta_{eff})]/2$, which is similar to the survival term a of SF14. The six components combine as:

$$S_{\text{mech}} = 0.15\, g_{\text{nuc}} + 0.10\, g_{\text{NB}} + 0.10\, g_{\text{BA}} + 0.10\, g_{\text{clamp}} + 0.25\, g_{\text{stab}} + 0.30\, g_{\text{angle}} \qquad \textbf{(SF1)}$$

**Šidák Multiplicity-Corrected Attack Angle:** For a carbon bearing $n$ equivalent scissile C-F bonds, the angular defect of the best-aligned bond is converted to a per-bond miss probability $p_1$, combined across the $n$ bonds by a Šidák survival term [4], and back-transformed to an effective angle so that a polyfluorinated carbon is not credited for the most favourable of $n$ independent chances (SF2). The graded angle component, $g_{angle}$, is evaluated with respect to $\theta_{eff}$. The correction is applied once, upstream of the score, so the tier scheme's angle criterion and $g_{angle}$ read the same corrected value. The corrected angle follows as:

$$d = \max(0, 180° - \theta); \quad p_1 = \frac{1 - \cos d}{2}; \quad \theta_{\text{eff}} = 180° - \arccos(1 - 2[1 - (1 - p_1)^n]) \qquad \textbf{(SF2)}$$

**Graded Chemical-Feasibility Factor:** A leaving-group term $f_{\mathrm{BDE}}$ falls linearly from one to zero as the scissile C-F bond-dissociation energy BDE rises across the 120 to 132 kcal mol$^{-1}$ window, following the established dependence of backside $S_N2$ leaving-group ability on C-F bond strength [5,6]. A β-withdrawal term $f_\beta$ discounts vicinal fluorination and ether oxygens ($n_\beta$), which withdraw electron density from the reactive centre [5] and track the substrate range of the characterised enzymes, which defluorinate difluoroacetate but not perfluorooctanoic acid [7,8]. Their product is floored at 0.10 (SF3), so a recalcitrant substrate is scaled down in rank but never excluded. The feasibility factor is defined as:

$$f_{\mathrm{BDE}} = \mathrm{clip}[\frac{132 - \mathrm{BDE}}{12}, 0, 1];$$
$$f_\beta = \frac{1}{1 + 0.35\, n_\beta}; \qquad \textbf{(SF3)}$$
$$\Phi = \max(0.10, f_{\mathrm{BDE}}\, f_\beta)$$

**Effective Score and Penalties:** The tier scheme and the elite gate both use the effective score (SF4), which is the geometric score less a graded chemistry penalty (SF5, with $O_{bs}$ the backside steric occlusion in Å) and a containment penalty, floored at zero. Feasibility does not enter here; it multiplies the competence score alone (SF14). Within the chemistry penalty, only the occlusion term fades as the trajectory approaches linearity, at full weight at or below 175° and zero at 180°, since a pose on the anti-periplanar axis has cleared the obstruction that term measures [6]. Bond strength and β-withdrawal are substrate properties and do not fade, and whose measurement failed is set to zero rather than imputed. An unresolved scissile centre zeroes the bond-strength term, and an unmeasured occlusion zeroes the occlusion term, and the complex is flagged and barred from the elite tier, so a mapping failure cannot invent a penalty or purchase a promotion. The effective score is formed as:

$$S_{\mathrm{eff}} = \max(0, S_{\mathrm{mech}} - P_{\mathrm{chem}} - P_{\mathrm{contain}}) \qquad \textbf{(SF4)}$$

$$P_{\mathrm{chem}} = 0.015 \max(0, \mathrm{BDE} - 123)$$
$$+ 0.13 \max(0, O_{\mathrm{bs}} - 2.0) \cdot s + 0.08\, n_\beta; \qquad \textbf{(SF5)}$$
$$s = \mathrm{clip}[\frac{180° - \theta_{\mathrm{eff}}}{5}, 0, 1]$$

**Near-Attack Conformation and Occupancy:** A trajectory frame qualifies as a near-attack conformation [3] when the catalytic triad is intact, both relay distances are within the SF1 gates, and the Oδ–Cα distance and the backside attack angle simultaneously satisfy the strict gates. The occupancy is the fraction of post-equilibration frames that qualify (SF6). Frames preceding equilibration are written to the per-frame record flagged as such and are excluded from both the numerator and the denominator, so the reported occupancy is conditioned on the equilibrated window rather than on the full trajectory. The frame criterion is stated as:

$$\begin{gathered}\mathrm{NAC}(t) = 1 \text{ if } d_{\mathrm{NB}}(t) \le 5.0\text{ Å and } d_{\mathrm{BA}}(t) \le 5.5\text{ Å} \\ \text{and } d_{\mathrm{OC}}(t) \le 3.2\text{ Å and } \theta(t) \ge 155°, \text{else } 0; \\ \mathrm{occupancy} = \frac{1}{T_{\mathrm{eq}}} \sum_{t \in \mathrm{post-eq}} \mathrm{NAC}(t)\end{gathered} \tag{SF6}$$

**Turnover (defluorination) Gate:** A lead is called defluorinating when the scissile C-F bond breaks measurably in the quantum mechanics/molecular mechanics (QM/MM) relaxed scan, with the product-end length reaching the geometric cleavage threshold (SF7) [1]. The departing-fluoride Mulliken charge [9], which under-reports the free-ion value in the diffuse-function-free basis, and the one-dimensional-scan barrier are reported as corroborating context, not as independent gates. The turnover gate is defined as:

$$\mathrm{Is}_{\mathrm{Defluorinating}} = 1 \text{ iff } d(\text{C-F})_{\mathrm{product}} \ge 1.8\text{ Å} \tag{SF7}$$

**Kruskal-Wallis Effect Size:** Cross-tier magnitude is quantified by the $\varepsilon^2$ statistic (SF8), the fraction of rank variance explained by the tier factor; $H$ is the Kruskal-Wallis statistic [10], $k$ is the number of tiers and $n$ is the total number of complexes. Observed: $\varepsilon^2 = 0.47$ (effective mechanistic score), 0.40 (mechanistic score), 0.26 (bimolecular nucleophilic substitution ($S_N2$) angle), 0.11 (nucleophile distance), all with $p$ returned as exactly zero by the analysis pipeline, which places the bound below the smallest representable double-precision value ($\approx 5 \times 10^{-324}$) rather than at any printed figure; $k = 7$, $n = 58{,}056$, the ranked set plus the six calibration controls. Test statistics, effect sizes and corrected p-values are reported in Supplementary Data SS_05. For the cross-tier Kruskal-Wallis tests, $\varepsilon^2$ is given in the effect_size column with effect_type = "epsilon$^2$", and $H$ is the test statistic, n_groups is the number of groups, and $n$ is the sample size. The effect size is computed as:

$$\varepsilon^2 = \frac{H - k + 1}{n - k} \tag{SF8}$$

**Interaction Affinity Score:** The protein-ligand interaction fingerprint is summarised as a weighted count of contact types minus an average-distance term (SF9). The geometric criterion for each contact type reproduces the Schrödinger Maestro default definitions and follows the primary literature for that interaction: hydrogen bonds [11], hydrophobic contacts [12], salt bridges [13,14], halogen bonds [15,16] and metal coordination [17]. The counted types are: hydrogen bonds (HB), hydrophobic contacts (HP), salt bridges (SB), halogen bonds (HAL), fluorine contacts (FL), fluorine-polar (FP), fluorous-hydrophobic (FH), fluorous-fluorous (FF) and metal coordination (MC). The weighted count is formed as:

$$A = 1.5\,\mathrm{HB} + 1.0\,\mathrm{HP} + 2.5\,\mathrm{SB} + 2.0\,\mathrm{HAL} + 1.0\,\mathrm{FL} + 4.0\,\mathrm{FP} + 2.0\,\mathrm{FH} + 3.0\,\mathrm{FF} + 2.5\,\mathrm{MC} - 0.05\,d_{\mathrm{avg}} \quad \textbf{(SF9)}$$

**WaterMap-Guided Quantum-Water Selection:** A candidate first-shell water is ranked by an effective distance in which a match to a thermodynamically stable WaterMap site (structural, $\Delta G < 0$) within 1.5 Å discounts the geometric distance (SF10) [18], so structural fluoride-stabilising waters are preferred for the quantum region over merely nearest ones. WaterMap sites are recorded in their own reference frame and are superimposed onto the selected molecular-dynamics frame by Cα Kabsch alignment before any distance is measured [19]. If fewer than three Cα atoms are shared, or the fold fit exceeds MD_FOLD_RMSD_MAX, the bias is dropped, and the selection reverts to nearest-first geometry; the same applies when no WaterMap data are available, in which case the discount is identically zero. The effective distance is evaluated as:

$$d_{\mathrm{eff}} = d - 1.0\max(0, -\Delta G_{\mathrm{site}}),\ \text{applied if } |\text{water} - \text{site}| \le 1.5\ \text{Å} \quad \textbf{(SF10)}$$

**Containment Penalty:** The effective score of SF4 subtracts a containment penalty that grows linearly as the ligand falls short of the target cavity containment (SF11). $C_{cav}$ is the fraction of ligand heavy atoms counted as cavity-contained, and containment at or above $CONTAIN\_PEN\_TARGET$ carries no penalty. Where the ray cast could not be completed, $C_{cav}$ is set to zero, and the pose therefore takes the full containment penalty, the opposite convention to the chemistry terms of SF5, which are zeroed on non-measurement. The asymmetry is deliberate: an unmeasured pose must not read as ideally buried and so escape the penalty, whereas an unmeasured bond strength must not manufacture one. The containment penalty is evaluated as:

$$P_{\mathrm{contain}} = 1.00\max(0, 0.85 - C_{\mathrm{cav}}) \quad \textbf{(SF11)}$$

**Cavity Containment:** A ligand heavy atom counts as cavity-contained once its buriedness reaches BURIAL_MIN (SF12). Buriedness is measured by casting BURIAL_RAYS rays per heavy atom on a Fibonacci sphere, each searched out to BURIAL_PROBE_A, with a ray counted as blocked when a protein heavy atom lies within BURIAL_RAY_CLEARANCE of the ray axis. The cavity fraction is computed as:

$$C_{\mathrm{cav}} = \frac{1}{N_{\mathrm{heavy}}} \sum_{i=1}^{N_{\mathrm{heavy}}} 1[b_i \geq 0.70] \qquad b_i = \frac{\text{blocked rays}}{42} \tag{SF12}$$

**Active-Site Containment:** The site8 term is the fraction of ligand heavy atoms lying within SITE8_SHELL_A of any active-site residue atom (SF13). It measures engagement with the reactive shell rather than fit to the cavity, so a tail leaving the shell sits outside the reactive site. The site8 fraction is computed as:

$$C_{\mathrm{site8}} = \frac{1}{N_{\mathrm{heavy}}} \sum_{i=1}^{N_{\mathrm{heavy}}} 1[\min_{j \in \mathrm{AS}} |r_i - r_j| \leq 5.0\ \text{Å}] \tag{SF13}$$

**Competence Score:** Tier_2A and Tier_2B are gated by a competence score built from six weighted components and clipped to the unit interval after multiplication by the feasibility factor (SF14), with floors of 0.50 and 0.40 respectively. The two elite tiers do not read it: they gate on geometry, the machinery gate and the assembly score. The same two floors are re-applied after those caps, so a geometry-strong but low-feasibility pose that is demoted into Tier_2A or Tier_2B, rather than entering it on the forward path, is dropped to Tier_3 and loses degrader status. Competence is also the first within-tier key in the mechanism-first ranking. Two details differ from SF1: the angle term is the raw Šidák survival [4] rather than the back-transformed effective angle of SF2, and Φ multiplies the competence score directly, whereas the effective score of SF4 carries no feasibility factor at all; chemical feasibility reaches the tier scheme only through the graded chemistry penalty of SF5. The competence score is formed as:

$$
\begin{aligned}
S_{\mathrm{comp}} &= \Phi[0.30\,a + 0.25\,\delta + 0.20\,\kappa + 0.10\,\tau + 0.10\,\rho + 0.05\,h] \\
a &= (1 - p_1)^n \\
\delta &= \mathrm{clip}[\frac{3.8 - d_{\mathrm{nuc}}}{3.8 - 2.4}, 0, 1] \\
\kappa &= \mathrm{clip}[\text{clamp integrity}, 0, 1] \\
\tau &= \mathrm{clip}[\frac{2.5 - \Delta_{\mathrm{traj}}}{2.5}, 0, 1] \\
\rho &= \frac{1}{2}(\mathrm{clip}[\frac{5.0 - d_{\mathrm{NB}}}{5.0 - 2.5}, 0, 1] + \mathrm{clip}[\frac{5.5 - d_{\mathrm{BA}}}{5.5 - 2.5}, 0, 1]) \\
h &= 1 \text{ if halide stabilised, else } 0
\end{aligned}
\qquad \textbf{(SF14)}
$$

**Catalytic Assembly Score:** The assembly score asks whether the eight-residue active site is assembled as in the crystal reference rather than merely present, based on the active-site root-mean-square deviation (RMSD) against 3R3U (SF15) [1]. It caps the attainable tier through the floors listed in Supplementary Table S1: a pose above zero but below every floor lands in Tier_4. B is set to zero, rather than evaluated, when the eight-residue superposition could not be measured, and it is that unmeasurable case which marks a decoy. The seven tiers are labelled Tier_1A through Tier_5_Decoy, matching the values in Supplementary Data SS_02. The assembly score is defined as:

$$
B = \frac{1}{1 + \mathrm{RMSD}_{\mathrm{AS}}} \qquad \textbf{(SF15)}
$$

**Rate-Weighted Ensemble Barrier:** Barriers from the scanned frames are combined by a Boltzmann-weighted estimator (SF16), which is dominated by the lowest barrier and is not a mean. Frames returning $Frame\_Barrier\_In\_Window = False$ resolved no transition state and are excluded before applying the estimator; the reaction energy is averaged with the same weights. Per-frame barriers, reaction energies and the in-window flag, together with the QM-region composition and net charge for every scanned system, are reported in Supplementary Data SS_14 [20,21]. N is the number of frames retained after exclusion, meaning those for which a barrier was parsed and an interior transition state resolved within the scan window, and $\Delta E‡\_min$ is the lowest barrier among those same retained frames. RT = 0.596 kcal $mol^{-1}$ at 300 K. N is reported per system as QSite_N_Frames_Scored, against QSite_N_Frames_Attempted for the number of frames scanned. The ensemble barrier is estimated as:

$$\Delta E^{\ddagger}_{\mathrm{ens}} = \Delta E^{\ddagger}_{\mathrm{min}} - RT\ln\left(\frac{1}{N}\sum_{i=1}^{N}\exp\left[-\frac{\Delta E^{\ddagger}_{i} - \Delta E^{\ddagger}_{\mathrm{min}}}{RT}\right]\right) \quad \textbf{(SF16)}$$

**Defluorination Propensity:** The propensity combines the strict near-attack probability with a Boltzmann factor in the barrier (SF17). It is a monotonic ranking proxy, not a rate: a rate constant requires the Gibbs barrier via the Eyring relation, and the pipeline runs no frequency calculation, so it carries no zero-point or thermal correction, no activation entropy and no prefactor [22]. Only the ordering is claimed. The propensity is undefined, rather than zero, when the scan resolved no transition state: a candidate whose barrier is unresolved is withheld from the ranking rather than placed at the bottom of it. A candidate with a resolved barrier but no strict near-attack population does have a genuine propensity of zero and is ranked last. $Defluor\_Propensity\_Norm$ rescales the quantity to the unit interval by min-max in log space, because the raw value spans many orders of magnitude. The propensity is computed as:

$$\mathrm{Propensity} = P(\text{strict-NAC}) \cdot \exp\left(-\frac{\Delta E^{\ddagger}}{RT}\right) \quad \textbf{(SF17)}$$

**Soft Catalytic Score:** A third, fully continuous score is computed alongside the mechanistic and competence scores, replacing every binary anchor of SF1 with a logistic function so that a hair's-breadth threshold crossing does not move a complex. Writing $\sigma(x;\, k, x_0) = 1/(1 + exp[-k(x - x_0)])$, the score combines nucleophile reach, attack angle and triad integrity with weights summing to one (SF18). The two distance terms carry negative steepnesses because they must fall with distance; the angle term rises. The score gates nothing and ranks nothing. It is reported as a fully continuous companion to SF1 and underlies the catalytic-geometry figures. The nucleophile and angle sigmoids are centred on the strict near-attack cutoffs they soften (NAC_DIST_STRICT 3.2 Å, NAC_ANGLE_STRICT 155°). The two triad sigmoids are centred on the triad-integrity thresholds rather than on the mechanistic-score relay gates: the Nuc-Base midpoint equals THRESHOLD_TRIAD_NB (4.5 Å), and the Base-Acid midpoint is set at 5.0 Å, below the 7.0 Å THRESHOLD_TRIAD_BA cutoff, so that the steepest scoring gradient falls in the 4–6 Å pre-reactive range; the hard cutoff remains the gate rather than the sigmoid centre. The soft score is formed as:

$$\begin{aligned} S_{\mathrm{soft}} &= w_{\mathrm{nuc}}\,\sigma(d_{\mathrm{nuc}};\, -4.0, 3.2\,\text{Å}) + w_{\mathrm{ang}}\,\sigma(\theta;\, 0.15, 155°) \\ &+ w_{\mathrm{int}}\,\sigma(d_{\mathrm{NB}};\, -2.0, 4.5\,\text{Å}) \cdot \sigma(d_{\mathrm{BA}};\, -2.0, 5.0\,\text{Å}) \quad \textbf{(SF18)} \\ w_{\mathrm{nuc}} &= 0.40, w_{\mathrm{ang}} = 0.30, w_{\mathrm{int}} = 0.30 \end{aligned}$$

Active-Site Conservation Score: The conservation score measures how well a candidate retains the catalytic machinery, active-site geometry, and sequence of the reference active site, and is available per complex in Supplementary Data SS_02. It combines three weighted components whose weights sum to one: machinery integrity (CONSERV_W_INTEGRITY), active-site geometric fit (CONSERV_W_GEO) and sequence identity (CONSERV_W_IDENT). The reference variant, reported as r3u_ActiveSite_Conservation_Score, drops the integrity term and reweights the remaining two, again summing to one. The conservation score is formed as:

$$C_{AS} = 0.60\, I_{mach} + 0.30\, G_{fit} + 0.10\, S_{ident};$$
$$C_{ref} = 0.25\, S_{ident} + 0.75\, G_{fit} \qquad \text{(SF19)}$$

***Supplementary Table S1 | Scoring and statistical constants.*** *Each constant below takes a single value and, except where a row states otherwise, is used unchanged in every computational step and figure. The conservation scores of SF19 are available per complex in Supplementary Data SS_02 as ActiveSite_Conservation_Score, r3u_ActiveSite_Conservation_Score and binding_likelihood_calc.*

| Symbol / constant | Value | Meaning |
|---|---|---|
| MECH_W_NUC / NB / BA | 0.15 / 0.10 / 0.10 | Mechanistic-score weights: nucleophile reach, nuc-base relay, base-acid relay |
| MECH_W_CLAMP / STAB / ANGLE | 0.10 / 0.25 / 0.30 | Clamp, halide stabilisation, Šidák-corrected attack angle [4] |
| MECH_CLASH_PENALTY | 0.03 | Subtracted per steric clash |
| MECH_CLASH_PENALTY_MAX | 0.15 | Cap on the total steric-clash penalty |
| MECH_NB_GATE / BA_GATE | 5.0 Å / 5.5 Å | Internal triad relay distance gates |
| MECH_ELITE_CONSTELLATION | 0.74 | Assembly value compensating for an intermediate mechanistic score for Tier_1A |
| MECH_ELITE_HI / LO | 0.90 / 0.85 | Effective-score thresholds in the Tier_1A machinery gate |
| COMP_W_ANGLE / DIST / CLAMP / TRAJ / TRIAD / HALIDE | 0.30 / 0.25 / 0.20 / 0.10 / 0.10 / 0.05 | Competence-score weights (SF14), sum 1.00 |
| SOFT_W_NUC / ANG / INT | 0.40 / 0.30 / 0.30 | Soft-score weights (SF18), sum 1.00 |
| SOFT_K_NUC / ANG / TRIAD | −4.0 / 0.15 / −2.0 | Logistic steepnesses; negative on the two distance terms so they fall with distance |
| SOFT_NB_MIDPOINT / BA_MIDPOINT | 4.5 Å / 5.0 Å | Relay sigmoid midpoints for the soft score, distinct from the SF1 relay gates |
| COMP_DIST_MIN / MAX | 2.4 Å / 3.8 Å | Nucleophile distances giving full and zero credit |
| COMP_TRAJ_MAX | 2.5 Å | Trajectory deviation giving zero credit |
| COMP_RELAY_MIN | 2.5 Å | Relay distance giving full triad credit |
| NAC_DIST_STRICT / ANGLE_STRICT | 3.2 Å / 155° | Strict near-attack gates (Oδ–Cα distance, attack angle) [3] |
| NAC_DIST_RELAXED / ANGLE_RELAXED | 3.8 Å / 145° | Relaxed near-attack gates [3] |
| SCISSILE_CF_BDE_MAX | 123 kcal mol$^{-1}$ | Scissile C-F BDE above which the bond is treated as too strong to cleave |
| SN2_BACKSIDE_OCCL_MAX | 2.0 Å | Backside steric-occlusion threshold for the $S_N2$ approach |
| FEAS_BDE_LO / HI | 120 / 132 kcal mol$^{-1}$ | Linear feasibility window for the leaving-group BDE term |

| | | |
|---|---|---|
| FEAS_BETA_PER_F | 0.35 | Per-β-withdrawal-unit feasibility discount (vicinal F or ether O) |
| FEASIBILITY_FLOOR | 0.10 | Lower bound on the feasibility factor |
| CHEM_PEN_W_BDE / OCCL / BETA | 0.015 / 0.13 / 0.08 | Graded chemistry-penalty weights (per unit BDE, occlusion, β) |
| CHEM_PEN_ANGLE_FULL / NONE | 175° / 180° | Angle window over which the occlusion penalty fades (SF5) |
| CONTAIN_PEN_TARGET / CONTAIN_PEN_W | 0.85 / 1.00 | Containment target and penalty weight (SF11) |
| BURIAL_RAYS / PROBE_A / RAY_CLEARANCE / MIN | 42 / 8.0 Å / 1.8 Å / 0.70 | Cavity-containment ray casting (SF12) |
| SITE8_SHELL_A | 5.0 Å | Active-site engagement shell (SF13) |
| TIER_ELITE_BDE_MAX | 128 kcal $mol^{-1}$ | Scissile C-F BDE ceiling for Tier_1A eligibility |
| TIER_1A_MAX_LIGAND_EXTENT | 5.0 Å | Ligand-extent value recorded for the controls-only validation subset; not a tier gate and not applied anywhere in the analysis, since the scheme is size-agnostic |
| TIER_CONSTELLATION_MIN | 0.55 / 0.45 / 0.35 / 0.25 | Assembly floors, Tier_1A to Tier_2B (SF15) |
| TIER_COMP_MIN | 0.50 / 0.40 | Competence-score floor at Tier_2A / Tier_2B; re-applied after the assembly and machinery caps |
| CONFLICT_CONF_HIGH / CONFLICT_CONF_LOW | 0.70 / 0.60 | Confidence thresholds for the conflict classification (main text); the two are deliberately asymmetric |
| TRUST_SCORE_EXCELLENT / ACCEPTABLE | 2.0 Å / 3.0 Å | Active-site superposition RMSD bands labelling control mapping trust as high, moderate or low |
| ALIGN_OPEN_GAP_SCORE / EXTEND | −10.0 / −0.5 | BLOSUM62 alignment penalties [23] |
| ALIGN_MIN_SEQ_IDENTITY | 25% | Mapping-reliability flag |
| ALIGN_GRADE_DEFS | A >90, B 80–90, C 70–80, D 60–70, E 50–60, F 40–50, G 30–40, H 20–30, I ≤20 (% identity) | Nine-grade banding of pairwise sequence identity, reported as Alignment_Grade; intervals are left-open and right-closed so a boundary value grades identically from both code paths |
| NUC_RESCUE_MAX_OFFSET_ELITE | 2 residues | Rescue offset cap for elite eligibility |
| QSITE_QM_WATER_RADIUS / MAX | 3.5 Å / 3 | First-shell quantum-water inclusion radius and count cap |
| QSITE_WM_WATER_MATCH_A / WEIGHT | 1.5 Å / 1.0 | WaterMap-site match radius and distance-discount weight [18] |
| MD_FOLD_RMSD_MAX | 3.0 Å | Cα fold-fit ceiling above which the WaterMap-site distance discount is dropped, and selection reverts to nearest-first (SF10) [18] |
| PHYS_MD_TIMESTEP_PS | (0.002, 0.002, 0.006) ps | RESPA inner, bonded and outer timesteps [24] |
| PHYS_MD_PRESSURE_BAR | 1.01325 bar | Production pressure (NPT) |
| PHYS_MD_THERMOSTAT_TAU / BAROSTAT_TAU | 1.0 ps / 2.0 ps | Temperature and pressure coupling times |
| PHYS_MD_NS / PHYS_MD_FRAMES | 1,000 ns / 100,000 | Production length and frames written, one every 10 ps; the frame pool from which the SF6 denominator is drawn |
| PHYS_MD_SEED | 2024 | Trajectory seed, distinct from the analysis seed of 42 |
| **ANALYSIS_SEED** | 42 | Seed for the analysis-side resampling and permutation procedures, distinct from the trajectory seed |

| | | |
|---|---|---|
| MD_EQUIL_BLOCKS / SKIP_FRAC / BLOCK_FRAC_MIN | 200 / 0.2 / 0.9 | Equilibration-detection blocking: number of blocks, leading fraction discarded, minimum fraction of subsequent blocks that must pass |
| MD_EQUIL_TARGET_T / T_TOL_K | 300.0 K / 3.0 K | Temperature target and tolerance for an equilibrated block |
| MD_EQUIL_V_TOL_PCT / V_DRIFT_MAX_PCT_NS | 1.0% / 0.05% $ns^{-1}$ | Box-volume tolerance and maximum drift |
| MD_EQUIL_CA_RMSD_MAX_A | 3.0 Å | Cα RMSD ceiling for an equilibrated block |
| CATION_CAP_DIST | 3.0 Å | Counter-ion approach to the nucleophile Oδ or substrate head group that disqualifies a frame as a QM/MM starting structure |
| POCKET_RESIDENCY_DIST | 8.0 Å | Nucleophile-distance cutoff defining a pocket-bound frame |
| QSITE_CF_CLEAVED_A | 1.8 Å | Product-end scissile C-F length at or beyond which cleavage is registered (SF7) [1] |
| QSITE_F_CHARGE_CLEAVED | −0.5 e | Product-state fluoride Mulliken-charge reference [9], reported as corroborating context (the cleavage call is geometric, SF7) |
| SCORE_W_AFF_HB / HP / SB / HAL / FL / FP / FH / FF / MC / DIST | 1.5 / 1.0 / 2.5 / 2.0 / 1.0 / 4.0 / 2.0 / 3.0 / 2.5 / 0.05 | Interaction-affinity weights (SF9) |
| BIND_INT_DENSITY_NORM / CROSS_PAE_NORM | 2.0 / 50.0 | F1 normalisers |
| BIND_LOGIT_CENTRE / GAIN | 2.5 / 1.25 | F2 centring and gain |
| BIND_LOGIT_CLAMP | 50.0 | Clamp on the logit before the sigmoid (F2); does not bind to any complex in this screen |
| GAS_CONSTANT_KCAL / MMGBSA_TEMPERATURE_K | $1.9872036 \times 10^{-3}$ / 300.0 | R (kcal $mol^{-1}$ $K^{-1}$) and T (K) for the rate-weighted ensemble estimator (SF16) |
| **Frame_Preorg_Score** | 100.0 / 5.0 / −50.0 / −10.0 | Pre-organisation weights per Å below the relaxed near-attack distance, per degree above the relaxed angle, per water-blockade unit and per kcal $mol^{-1}$ of unfavourable WaterMap ΔG. Applied as an eligibility mask on the scanned frames rather than as the ranking key; the two-stage selection rule is stated in the main-text QM/MM Methods. Values are available per frame in Supplementary Data SS_14. |

### Supplementary Figures (Results):

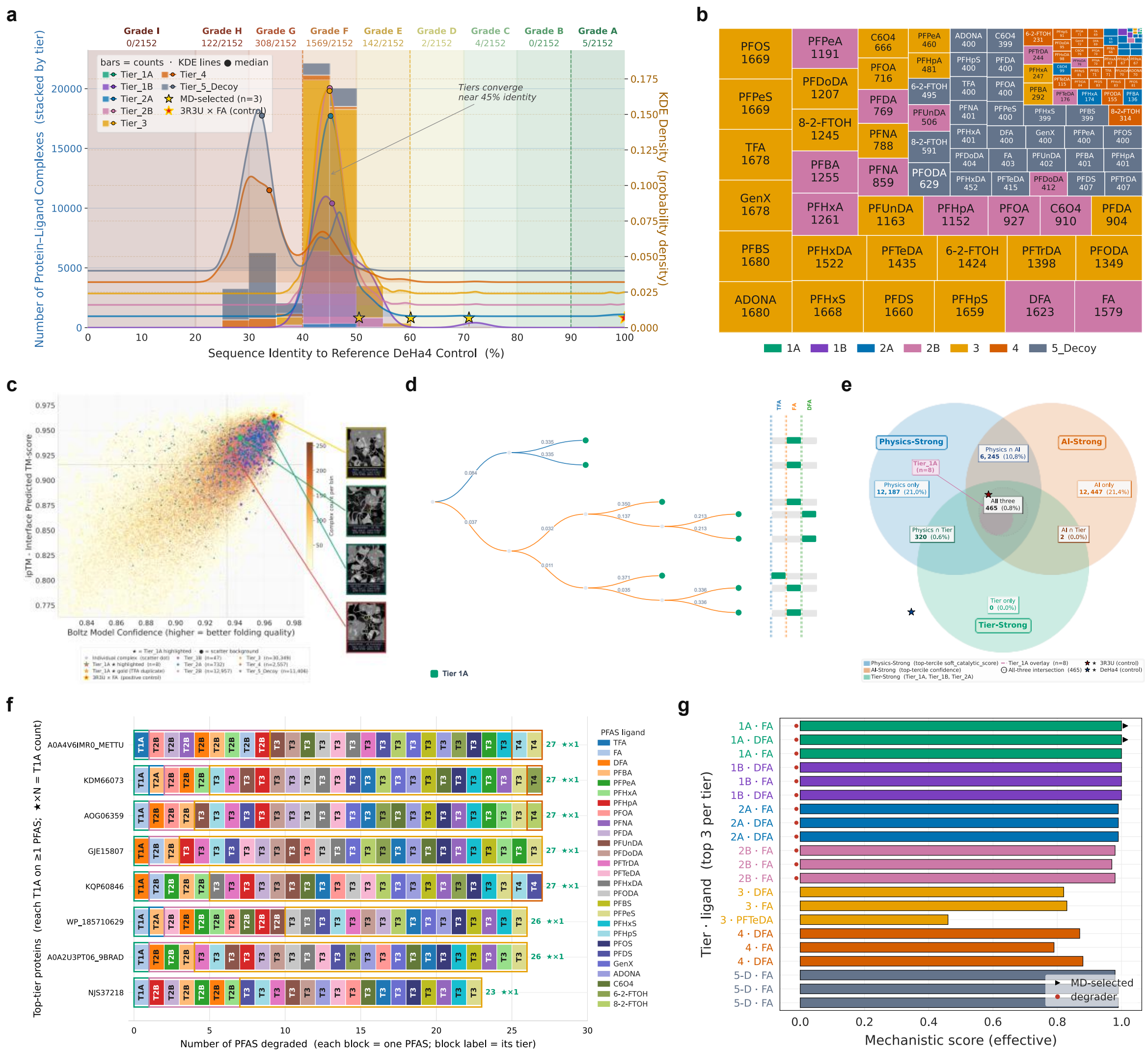


***Supplementary Figure 1 | Screening and scope diagnostics. Further characterisation of the input library and the PFAS scope. (a)*** *Distribution of active-site alignment grades across the library.* ***(b)*** *Treemap of the tier-by-ligand composition of the screened complexes.* ***(c)*** *AI-quality feature space (pTM, ipTM and confidence).* ***(d)*** *Tier_1A enzyme-by-PFAS dendrogram and cross-ligand tier matrix for fluoroacetate, trifluoroacetate and difluoroacetate.* ***(e)*** *Euler diagram of substrate-category overlap.* ***(f)*** *Top-tier protein-by-PFAS breakdown.* ***(g)*** *Swimmer plot of the top hits per tier.*

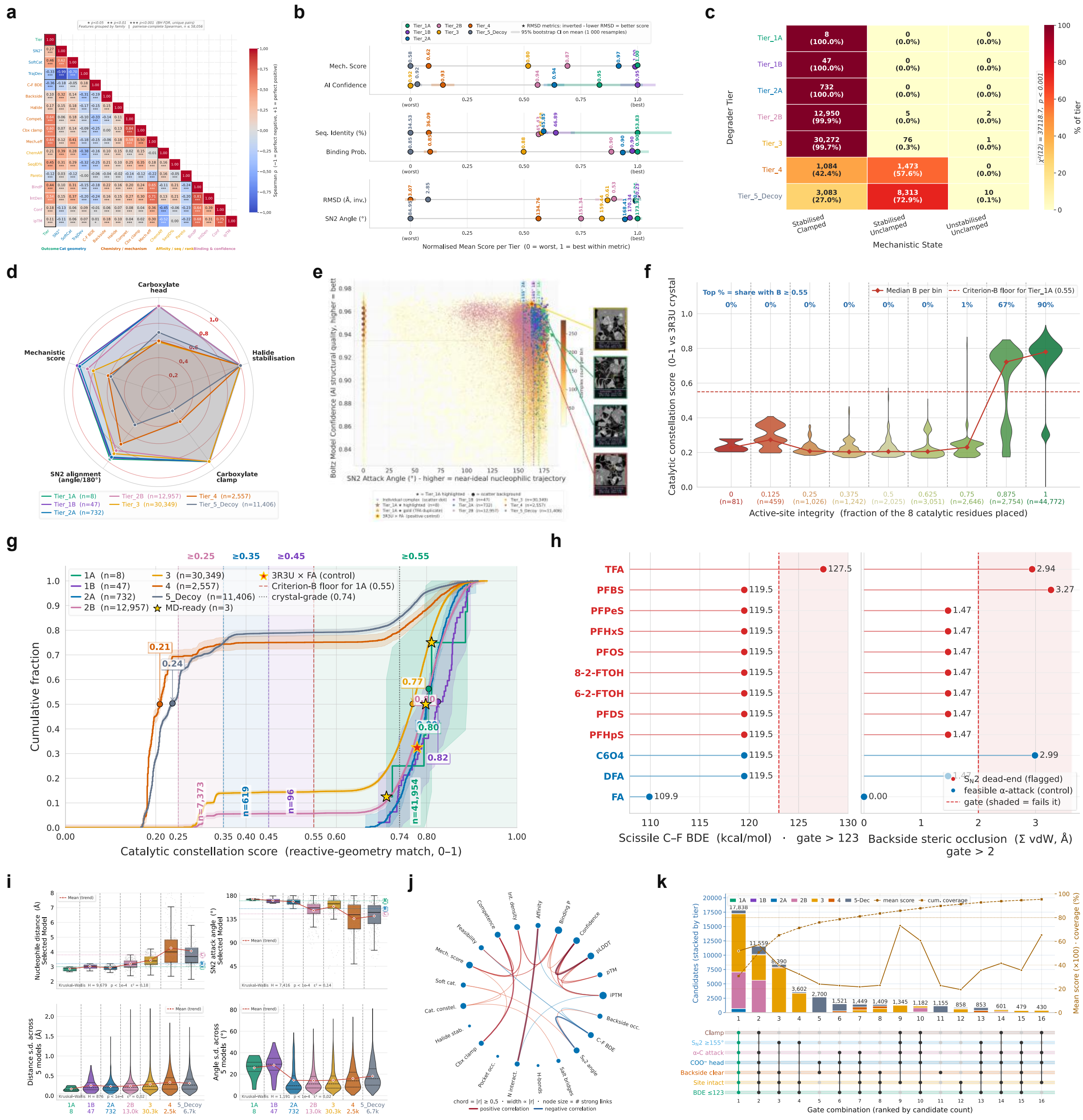


**Supplementary Figure 2 | Catalytic-mechanism diagnostics.** *Analysis of the mechanism-first score that expands Figure 4.* ***(a)*** *Pairwise correlations among the mechanistic features.* ***(b)*** *Tier-quality dot plot.* ***(c)*** *Cross-tabulation of discrete mechanistic states.* ***(d)*** *Mechanistic fingerprint of the tiers.* ***(e)*** *Mechanistic-quality feature space.* ***(f)*** *Criterion A gating criterion B.* ***(g)*** *Empirical cumulative distribution of criterion B by tier.* ***(h)*** *$S_N2$ dead-end gate.* ***(i)*** *Joint geometry-and-uncertainty view.* ***(j)*** *Metric co-variation network.* ***(k)*** *Mechanistic-gate intersection (UpSet) analysis, showing that 17,838 complexes (30.7%) satisfy every gate simultaneously, and that clearing every gate does not by itself confer elite status.*

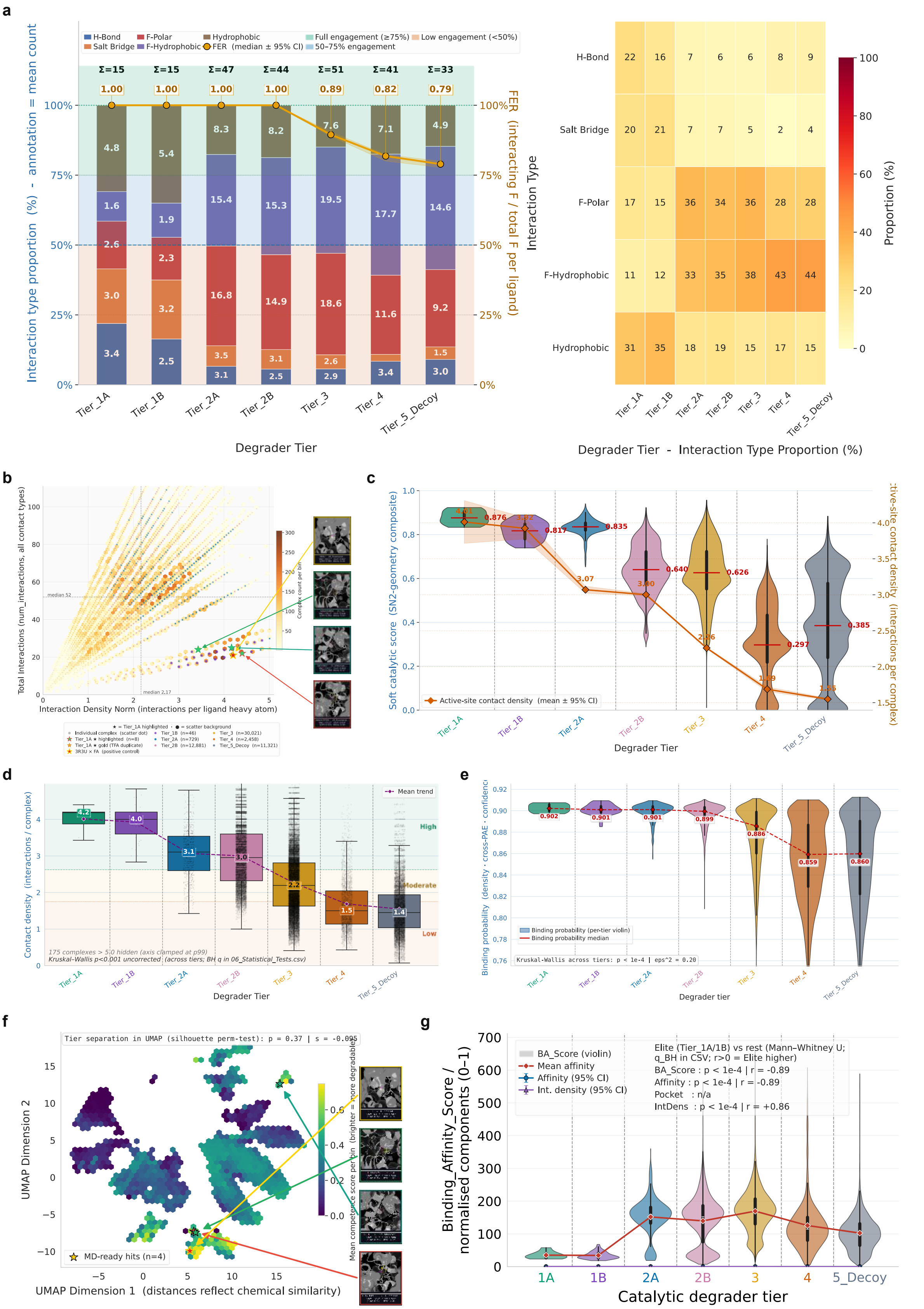


***Supplementary Figure 3 | Ligand interactions and chemical space.*** *Expanded interaction and chemical-space analysis supporting Figure 4.* ***(a)*** *Molecular interaction profile.* ***(b)*** *Interaction-quality*

*feature space. **(c)** Catalytic quality against predicted inhibition, separating productive from non-productive binding. **(d)** Active-site contact density by tier. **(e)** Binding energetics. **(f)** Chemical-space map. **(g)** Binding-affinity metric distributions across the tiers.*

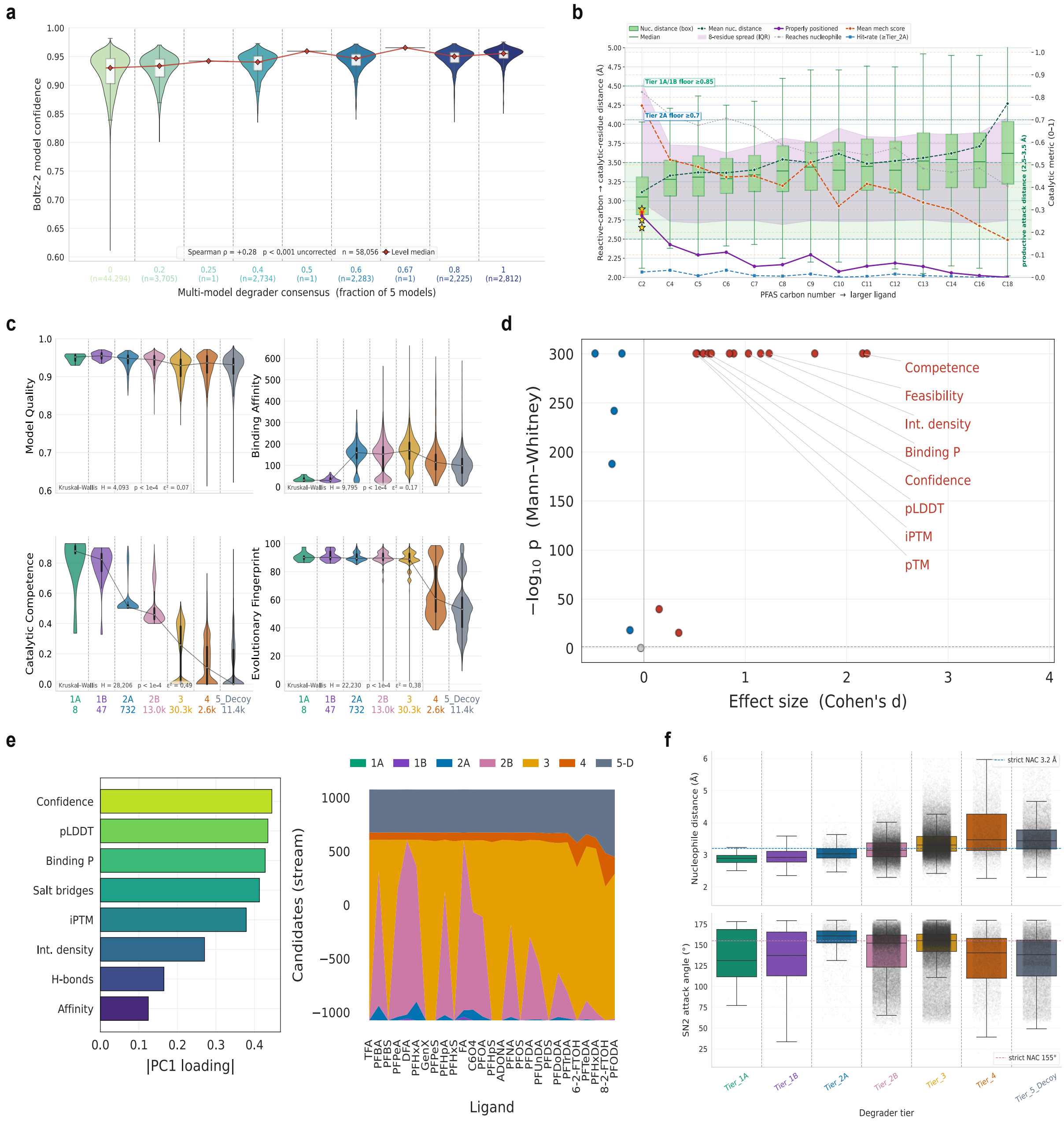

***Supplementary Figure 4 | Multi-model and diagnostic trends.** Diagnostic trends across the screen and the robustness of the ranking across the Boltz-2 model ensemble. **(a)** Confidence against consensus. **(b)** Reactive engagement. **(c)** Pillar divergence by tier. **(d)** Volcano plot of metric significance. **(e)** Feature-importance tier streams. **(f)** Multi-model geometry variance for the ipTM-90 subset.*

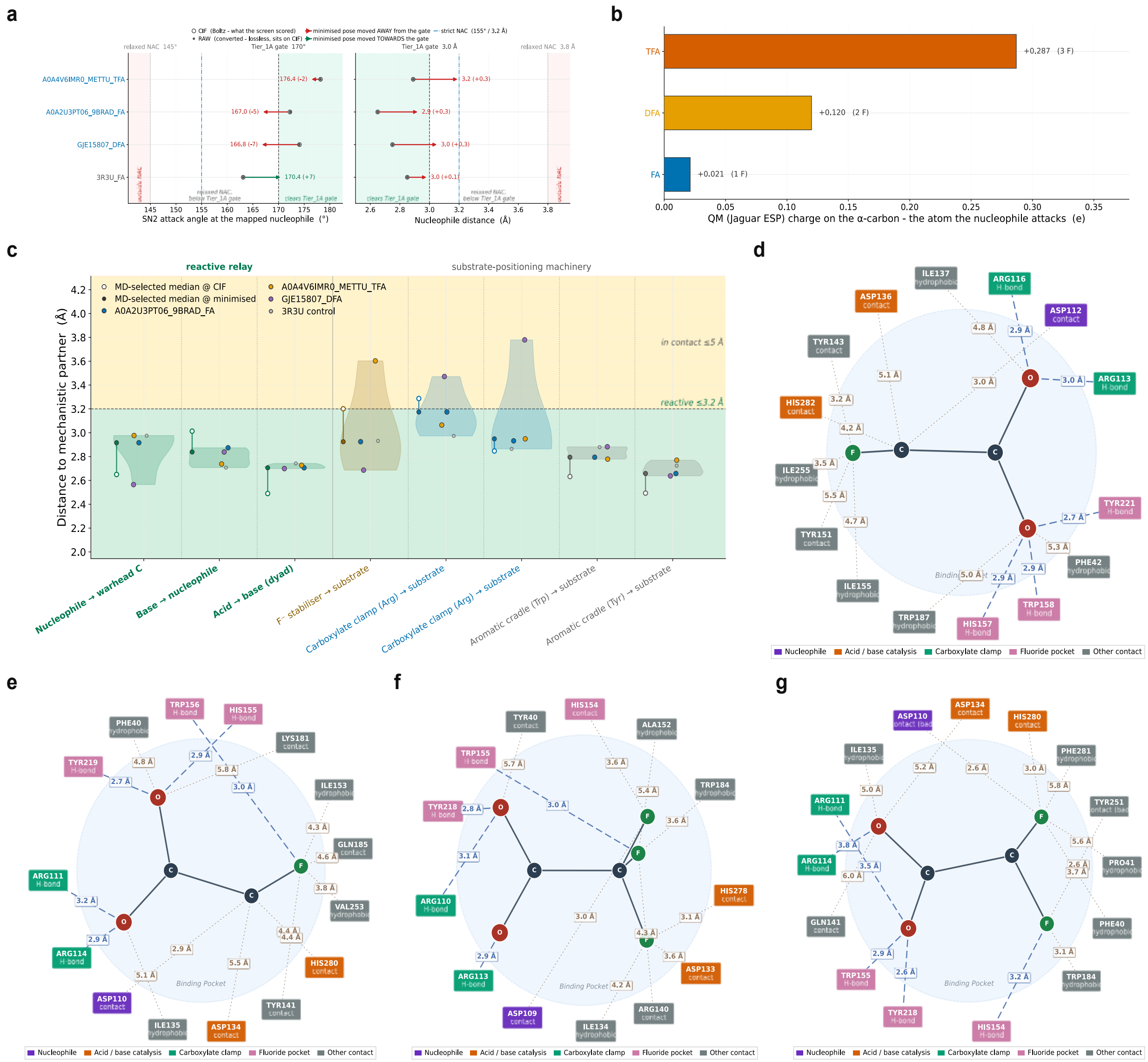


**Supplementary Figure 5 | Preparation, ESP and interaction maps.** *Preparation quality control for the systems advanced to physics validation.* ***(a)*** *Pose drift from the co-fold CIF to the prepared structure.* ***(b)*** *Electrostatic-potential-fitted α-carbon charge of the reactive ligands.* ***(c)*** *Distribution of machinery engagement.* ***(d–g)*** *Prepared-complex protein-ligand interaction maps, in the order of the 3R3U control, R1 (the fluoroacetate lead), R8 (the trifluoroacetate lead) and R2 (the difluoroacetate lead).*

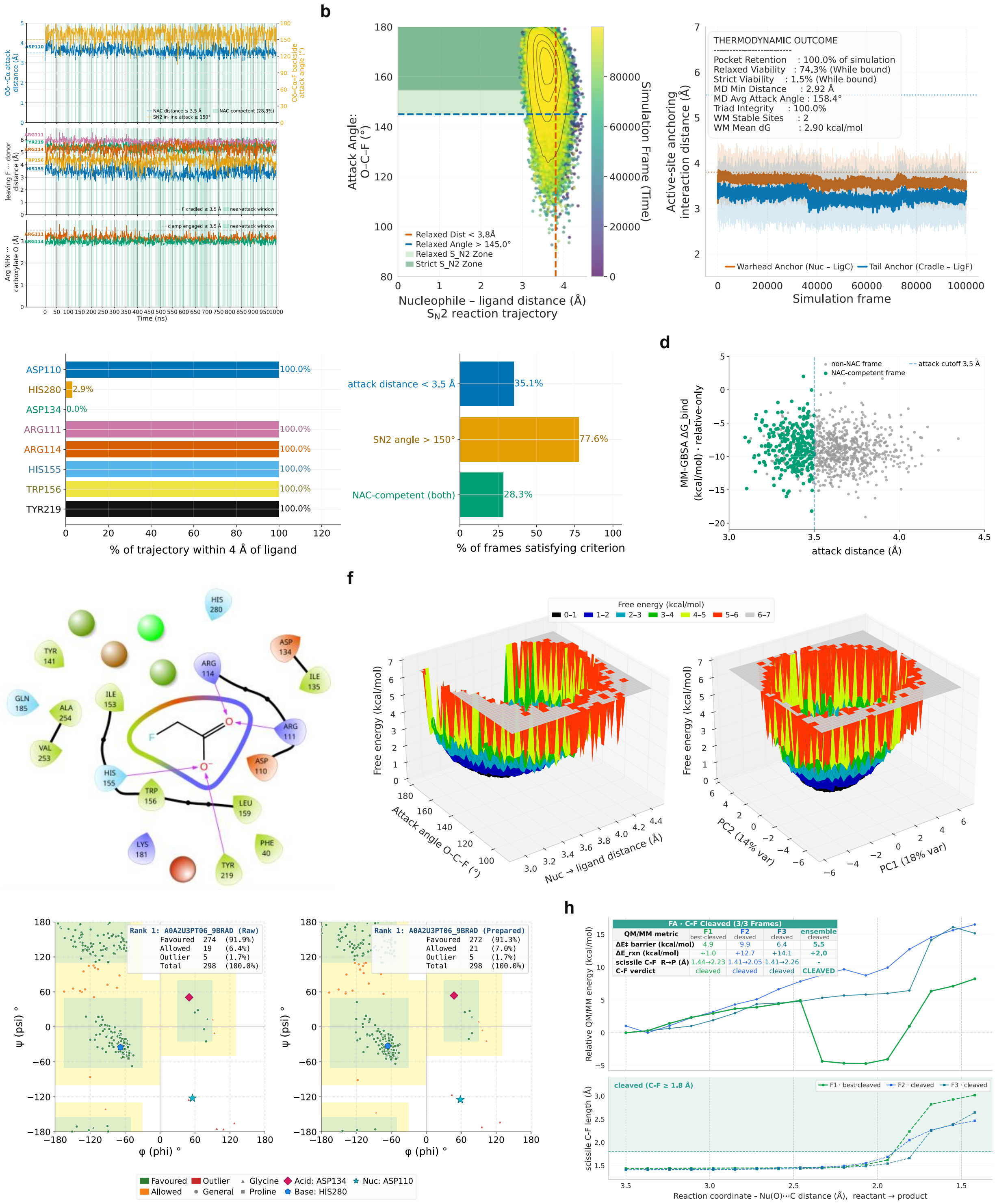


***Supplementary Figure 6 | Near-attack dynamics and C-F cleavage, R1.*** *Per-system near-attack dynamics and the C-F cleavage reaction for R1, the fluoroacetate lead (Bradyrhizobium sp., UniProt A0A2U3PT06), over the one-microsecond trajectory.* ***(a)*** *Reactive-pose trajectory (attack angle, halide-cradle and carboxylate-clamp distances over time).* ***(b)*** *Near-attack-conformation dashboard (attack-angle-versus-distance basin and active-site anchoring).* ***(c)*** *Per-residue occupancy and NAC-criteria summary.* ***(d)*** *Binding (MM-GBSA) against reactivity, per frame.* ***(e)*** *WaterMap hydration sites for this*

*system.* ***(f)*** *Free-energy landscapes of the reaction coordinate.* ***(g)*** *Prepared-pose comparative Ramachandran plot (reference against prepared structure).* ***(h)*** *QM/MM relaxed-scan reaction profile over the scissile C-F bond: the upper plot is the scan energy of the three frames with the per-frame metric table, and the lower plot is the scissile C-F length against the 1.8 Å cleavage band and found that all three frames cleave. The scan drives only the forming nucleophile-carbon coordinate, so the C-F elongation is in part a mechanical consequence of that drive and is shared by the reference (Supplementary Figure 9h); the panel therefore demonstrates geometric feasibility rather than a ranking, a measure of competence or a demonstration of turnover. Protein and ligand RMSD, RMSF and temperature are in Figure 6b; the ligand-protein 2D interaction map in Figure 6f–i; cross-rank MM-GBSA and machinery are in Figure 6d and Figure 8. Source data: SS_14.*

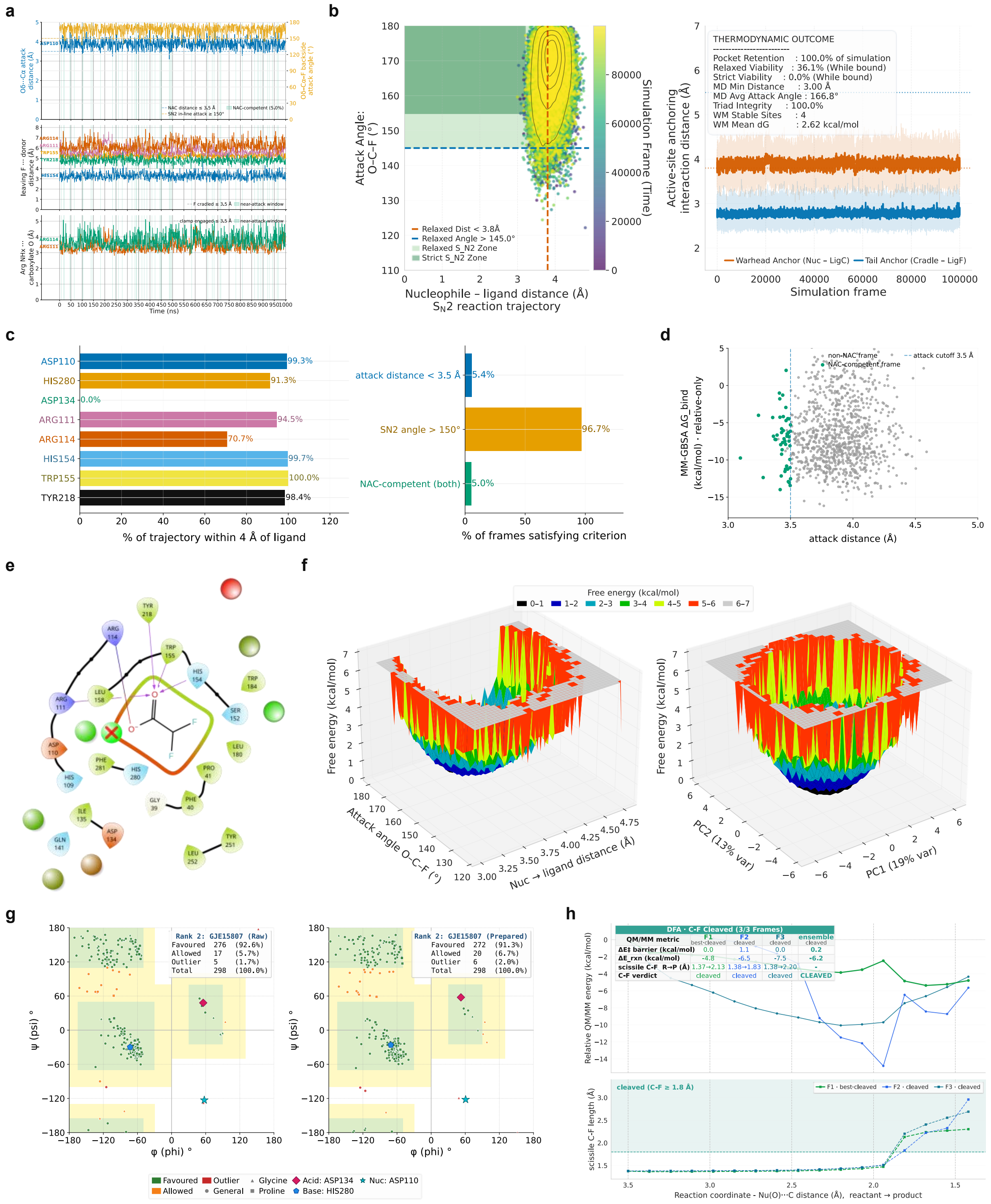


**Supplementary Figure 7 | Near-attack dynamics and C-F cleavage, R2.** *Per-system near-attack dynamics and the C-F cleavage reaction for R2, the difluoroacetate lead (Methylobacterium marchantiae, GenBank GJE15807), over the one-microsecond trajectory.* ***(a)*** *Reactive-pose trajectory (attack angle, halide-cradle and carboxylate-clamp distances over time).* ***(b)*** *Near-attack-conformation dashboard (attack-angle-versus-distance basin and active-site anchoring).* ***(c)*** *Per-residue occupancy and NAC-criteria summary.* ***(d)*** *Binding (MM-GBSA) against reactivity, per frame.* ***(e)*** *WaterMap hydration sites for this system.* ***(f)*** *Free-energy landscapes of the reaction coordinate.* ***(g)*** *Prepared-pose comparative Ramachandran plot (reference against prepared structure).* ***(h)*** *QM/MM relaxed-*

*scan reaction profile over the scissile C-F bond: the upper plot is the scan energy of the three frames with the per-frame metric table, and the lower plot is the scissile C-F length against the 1.8 Å cleavage band, and we found that all three frames cleave. The scan drives only the forming nucleophile-carbon coordinate, so the C-F elongation is in part a mechanical consequence of that drive and is shared by the reference (Supplementary Figure 9h); the panel therefore demonstrates geometric feasibility rather than a ranking, a measure of competence or a demonstration of turnover. Protein and ligand RMSD, RMSF and temperature are in Figure 6b; the ligand-protein 2D interaction map in Figure 6f–i; cross-rank MM-GBSA and machinery are in Figure 6d and Figure 8. Source data: SS_14.*

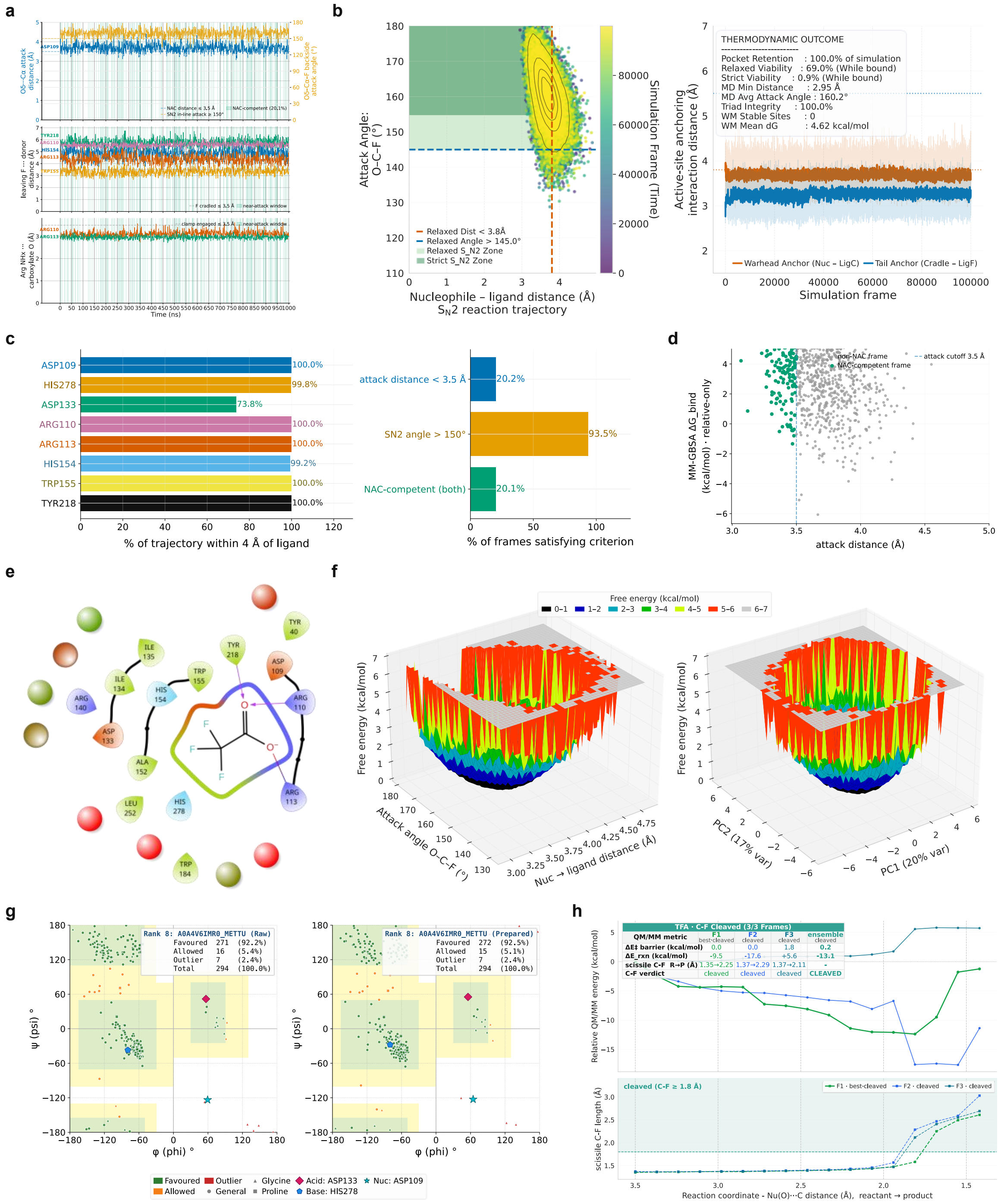

***Supplementary Figure 8 | Near-attack dynamics and C-F cleavage, R8.*** *Per-system near-attack dynamics and the C-F cleavage reaction for R8, the trifluoroacetate lead (Methylocella tundrae, UniProt A0A4V6IMR0), over the one-microsecond trajectory.* ***(a)*** *Reactive-pose trajectory (attack angle, halide-cradle and carboxylate-clamp distances over time).* ***(b)*** *Near-attack-conformation dashboard (attack-angle-versus-distance basin and active-site anchoring).* ***(c)*** *Per-residue occupancy and NAC-criteria summary.* ***(d)*** *Binding (MM-GBSA) against reactivity, per frame.* ***(e)*** *WaterMap hydration sites for this system.* ***(f)*** *Free-energy landscapes of the reaction coordinate.* ***(g)*** *Prepared-pose comparative Ramachandran plot (reference against prepared structure).* ***(h)*** *QM/MM relaxed-scan reaction profile over the scissile C-F bond: the upper plot is the scan energy of the three frames with the per-frame metric table, and the lower plot is the scissile C-F length against the 1.8 Å cleavage band and all three frames cleave. The scan drives only the forming nucleophile-carbon coordinate, so the C-F elongation is in part a mechanical consequence of that drive and is shared by the reference (Supplementary Figure 9h); the panel therefore demonstrates geometric feasibility rather than a ranking, a measure of competence or a demonstration of turnover. Protein and ligand RMSD, RMSF and temperature are in Figure 6b; the ligand-protein 2D interaction map in Figure 6f–i; cross-rank MM-GBSA and machinery are in Figure 6d and Figure 8. Source data: SS_14.*

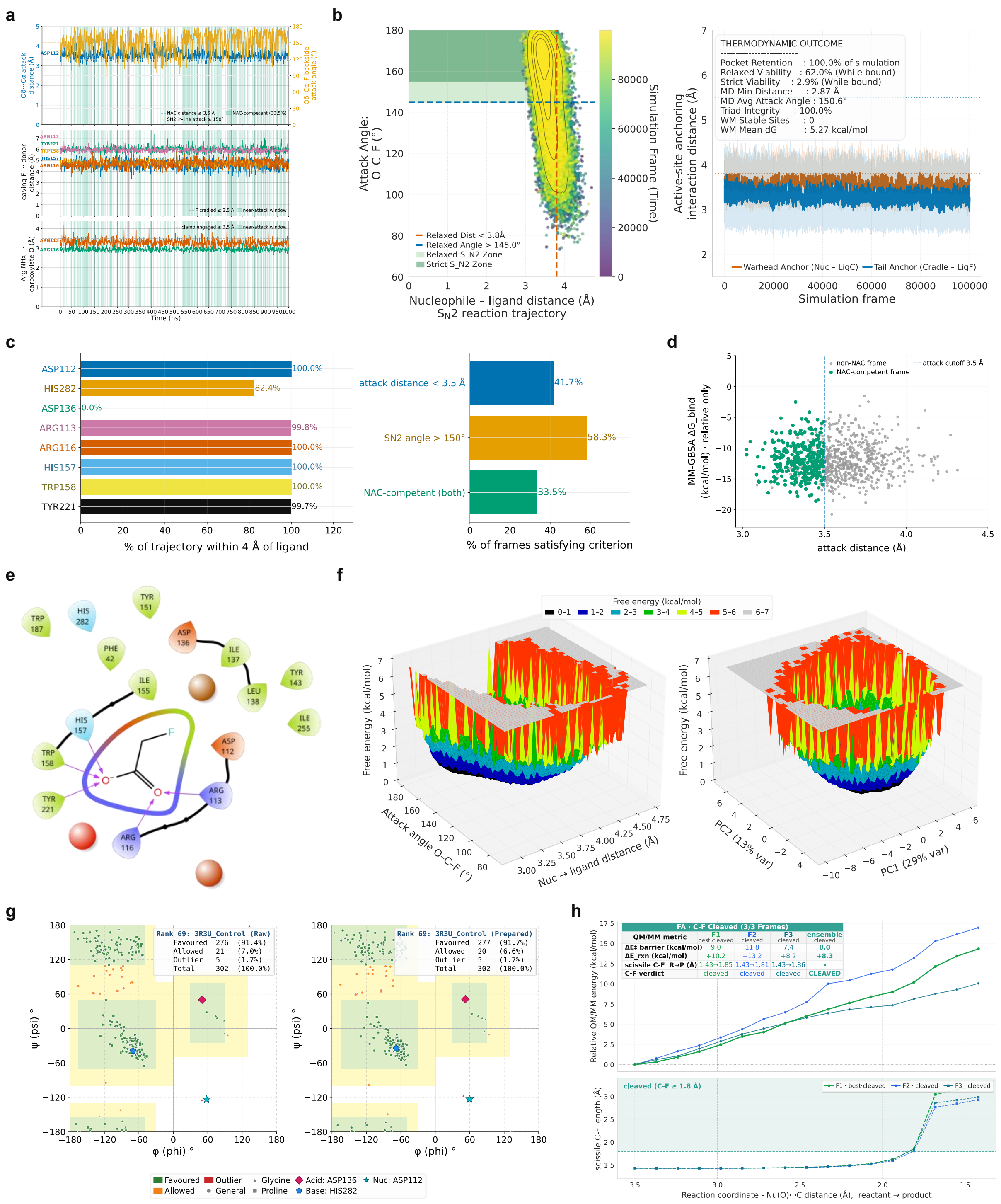


***Supplementary Figure 9 | Near-attack dynamics and C-F cleavage, R69.*** *Per-system near-attack dynamics and the C-F cleavage reaction for R69, the RPA1163/3R3U reference on its native substrate fluoroacetate, over the one-microsecond trajectory.* ***(a)*** *Reactive-pose trajectory (attack angle, halide-cradle and carboxylate-clamp distances over time).* ***(b)*** *Near-attack-conformation dashboard (attack-angle-versus-distance basin and active-site anchoring).* ***(c)*** *Per-residue occupancy and NAC-criteria summary.* ***(d)*** *Binding (MM-GBSA) against reactivity, per frame.* ***(e)*** *WaterMap hydration sites for this system.* ***(f)*** *Free-energy landscapes of the reaction coordinate.* ***(g)*** *Prepared-pose comparative*

*Ramachandran plot (reference against prepared structure).* ***(h)*** *QM/MM relaxed-scan reaction profile over the scissile C-F bond: the upper plot is the scan energy of the three frames with the per-frame metric table, and the lower plot is the scissile C-F length against the 1.8 Å cleavage band, and all three frames cleave. The scan drives only the forming nucleophile-carbon coordinate, so C-F elongation is partly a mechanical consequence of that drive and is shared by this reference system; the panel therefore demonstrates geometric feasibility rather than ranking, competence, or turnover. Protein and ligand RMSD, RMSF and temperature are in Figure 6b; the ligand-protein 2D interaction map is in Figure 6f–i; cross-rank MM-GBSA and machinery are in Figure 6d and Figure 8. Source data: SS_14. Residues are numbered as in the prepared model, which runs two positions higher than the deposited 3R3U numbering used elsewhere in this work (Asp112 here corresponds to the canonical Asp110).*

---

#### Declarations

**Ethical responsibilities:** not applicable; the study is computational.

**Consent for publication:** all authors consent to submission.

**Funding:** This work was supported by a Sapere Aude starting grant "Solving Microbial Degradation of PFAS" from the Independent Research Fund Denmark (DFF; grant number 5254-00090B).

**Acknowledgement:** The authors thank the members of the Microbial Ecology and Biotechnology Section (Now Environmental Sciences).

**Data Availability:** The data underlying all figures and tables are provided in the supplementary workbook Supplementary_Sheets_DeFluorX.xlsx, which is permanently archived on Zenodo (https://doi.org/10.5281/zenodo.23318610). The workbook also includes the source protein sequences and ligand structures used in this study. The Index sheet provides a mapping of each SS_nn identifier to its corresponding worksheet and contents.

**Code Availability:** The DeFluorX pipeline code is available at https://github.com/KU-MGB/DeFluorX.

**Competing interests:** The authors declare no competing interests.

**Author Contributions:** **SA** designed and implemented the pipeline, performed the analysis, wrote all the code, and wrote and revised the manuscript. **AF** contributed to the study design and provided experimental insight that informed pipeline development. **BT** contributed to methodology design and optimisation. **JBR** contributed to methodology design and optimisation. **MO** contributed to the study design and provided experimental insight that informed pipeline development. **TKN** secured funding, supervised the study, contributed to the study design, and critically revised the manuscript. All authors reviewed and approved the final manuscript.
